# Genome-wide fixation under viability selection

**DOI:** 10.1101/2022.09.19.508488

**Authors:** Bjarki Eldon

## Abstract

We investigate through simulations the effect of random sweepstakes, recurrent bottlenecks, dominance, and epistasis on selection a diploid population when the phenotype is determined by the genotype at a number of unlinked sites. We assume there are two types at each site, the wild type and a type conferring selective advantage. The focus is on identifying main trends in how the stated elements impact selection when selection acts on at least two unlinked site. We give examples of fixation trajectories under an array of scenarios, and record corresponding numerical estimates of the probability of fixation of the fit type at all the sites in question each time, and the time to fixation conditional on fixation. The results suggest that joint fixation of the fit type at the sites can occur in a number of scenarios, despite starting with the fit type in a single copy at each site. Random sweepstakes generally reduce both the probability and the time to fixation (conditional on fixation). Recurrent strong bottlenecks generally reduce the probability of fixation, but the effect of bottlenecks on the time to fixation (conditional on fixation), depends on the dominance mechanism in question. We identify an example of epistasis where fixation can occur, when in the absence of epistasis fixation does not occur at all or with much lower probability. Our results, coupled with an extensive analysis of genomic data of Atlantic cod(4), is further evidence for pervasive strong positive selection as being a key driving force in the evolution at least of highly fecund populations.

## 1 Introduction

Natural selection is a fundamental element shaping the evolution of populations. Understanding how selection interacts with other aspects of the biology of natural populations, such as recruitment dynamics, demography, epistasis, and dominance mechanisms is therefore essential. Single-locus models of selection have been much investigated. However, models of selection involving at least two (linked) loci or sites, polygenic selection, are increasingly being considered(8; 19; 66; 67), and for good reason. Genome-wide association studies in humans suggest that many phenotypes are influenced by a significant number of genetic variants(64). Molecular evidence of selective sweeps and polygenic adaptation also exist(18).

Here we investigate how epistasis among unlinked sites affects selection acting on the sites. Initially epistasis was used to refer to the masking of the effects of genetic variants(9). Epistasis may be prevalent and pervasive(49), play a significant role in molecular evolution(7; 15; 52; 60; 65; 69; 76), has been worked into the infinitesimal model(8), and may hold a key to understanding missing heritability(39). Nevertheless, little is known about the molecular mechanisms of epistasis(48). Fisher’s geometric model(31) is often taken as a reference model in empirical studies of epistasis(33), however it may not adequately explain the complete structure of fitness landscapes(14). Epistasis may play an important role in a number of aspects of biological populations such as diploidy, dominance mechanisms, and genome complexity(62). Further, epistasic interactions between more than two sites (higher order epistasis) may be common in nature (70; 71). Here we take epistasis to mean the dependence of the dominance mechanism at one site on the genotypes at other site(s), or the ‘genetic background’ of the given site.

Dominance refers to how homozygous and heterozygous genotypes at a site in a diploid population affect a phenotype, and is a key concept in molecular genetics(40), and therefore also in evolutionary genetics. Essentially it is about what the heterozygote (an individual carrying both types at a given site) does, or the phenotype of the heterozygote in relation to the phenotype of the two homozygotes (individuals carrying only one of two types at a given site). The notion of dominance can be traced at least to R.A. Fisher(30), S. Wright(74), and J.B.S. Haldane(35). It has been argued that most new mutations arise as recessive to the wild type (the type present in the population at the time of mutation to a new type)(72). For example, molecular studies of the bacteriophage *φ*6 yielded additive deleterious mutations and beneficial recessive mutations(41). Another type of dominance, overdominance (where the heterozygote is fitter than both homozygotes) plays a key role in maintaining genetic diversity(1; 51). However, the dominance mechanism of most new mutations that confer advantage in natural populations is not well known(57).

Here we use simulations to investigate the impact of random sweepstakes (a particular mechanism of recruitment dynamics)(4), recurrent bottlenecks, epistasis and dominance mechanisms on viability selection when the phenotype is a function of the genotype at a number (at least two) of unlinked sites in a diploid population. A ‘highly fecund’ population is one in which individuals have the potential to produce at least on the order of the population size number of potential offspring (juveniles) at any given time. Consider a highly fecund population characterized by broadcast spawning and Type III survivorship. Random sweepstakes is the chance matching of reproduction with favorable environmental conditions, thus providing a mechanism for skewed individual recruitment success (offspring number distribution), or sweepstakes reproduction. Another mechanism generating sweepstakes reproduction in highly fecund populations involves natural selection(73), and has been referred to as selective sweepstakes(4). Random and selective sweepstakes are two mechanisms turning high fecundity into skewed individual recruitment success. Sweepstakes reproduction has been suggested to be a significant factor in shaping genetic diversity among highly fecund marine broadcast spawners(2; 37; 38). Comparison of models of random sweepstakes(63) with actual population genetic data suggests that models of random sweepstakes may have an important role in explaining genetic diversity in such populations(3; 10; 13). An extensive analysis using whole-genome DNA sequence data suggests instead that recurrent and pervasive selective sweeps of strongly beneficial types arising by mutation(22) are at least as relevant as random sweepstakes in explaining genetic diversity in the highly fecund Atlantic cod(4). The simultaneous presence of abundant genetic variation and pervasive strong positive selection, as observed in Atlantic cod(4), suggests the presence of significant multivariate genetic constraints(69), implying widespread and epistasis between many sites.

Models of the evolution of genetic type frequencies under random sweepstakes are in the domain of attraction of a jump diffusion(11). Take 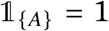 if *A* holds, and zero otherwise. Denote by {*B*_*t*_; *t* ≥ 0} standard Brownian motion, *E* = (0, *t*] × (0, 1] × [0, 1], *N* a Poisson process on [0, ∞) × (0, 1] × [0, 1] with intensity measure d*t* ⊗ *r*^−2^*F*(d*r*) ⊗ d*u* where *F* is a finite measure on the unit interval without an atom at zero and a fraction *r* is replaced at rate *r*^−2^*F* (d*r*) in a haploid population evolving according to random sweepstakes. Let {*X*_*t*_ } ≡ {*X*_*t*_; *t* ≥ 0} denote a process tracking type frequency in a haploid population. Under random sweepstakes {*X*_*t*_ } is the solution of the SDE

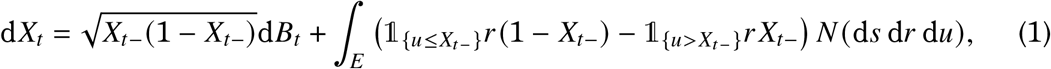

see (6; 11) for further details. It is thus clear that methods based on continuous diffusion(44; 45) do not apply to models of random sweepstakes. Mathematical analysis of models of random sweepstakes with selection is therefore a bit involved. Published work in this direction is, as far as we know, restricted to the haploid single-locus framework(6; 20; 21; 27; 32).

Incorporating recurrent bottlenecks corresponds to studying genome-wide selection under random sweepstakes in a changing environment. Previous studies on the establishment of a new mutation in a varying environment have focused on a single site in a population without random sweepstakes and where the fluctuations in the environment are small(58). Here we will be concerned with strong bottlenecks, i.e. when the population size drops significantly below its starting value. Strong bottlenecks can be shown to significantly alter the excursion of a type frequency in a population evolving according to the Wright-Fisher model, i.e. leading to Ξ-Fleming-Viot processes(12). One might therefore expect that strong recurrent bottlenecks would have an effect on selection.

Simulations in the single locus framework have shown that random sweepstakes (without bottlenecks) lower both the the probability of fixation and the expected time to fixation conditional on fixation of the type conferring selective advantage at a single locus in a haploid population evolving according to random sweepstakes as in Eq (2) (adapted to haploidy) but without randomizing on *α*(25). Similar results (also based on simulations) have been obtained for a single site in a diploid population experiencing random sweepstakes, recurrent bottlenecks, and various dominance mechanisms using the model in Eq (2) and randomizing on *α*(26). Here we extend these investigations to the case of two or more unlinked sites, which allows to investigate the joint fixation at several unlinked sites of types conferring advantage, and the effects of varying dominance mechanisms among the sites and epistasis on selection.

## 2 The model

Our formulation builds on the Schweinsberg model of a haploid population evolving according to random sweepstakes(63). Consider a diploid population evolving in an environment with a fixed carrying capacity *C*, i.e. *N*_*t*_ ≤ *C* for all *t* ≥ 0 where *N*_*t*_ is the population size at time (generation) *t*, or the number of diploid individuals in the population at time *t*. In any given generation, the current individuals randomly form pairs, and each pair independently contributes a random number of juveniles, or potential offspring. After producing juveniles all the parents perish. If *N*_*t*_ is odd we assume ⌊*N*_*t*_/2⌋ pairs produce juveniles, and one random individual will be left out. Let *X* denote the random number of juveniles contributed by a given parent pair, with

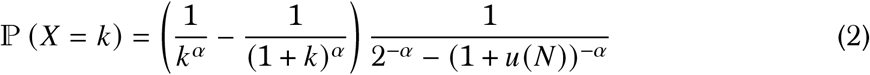

for *k* ∈ {2, 3, …, *u*(*N*)}, where *u*(*N*) is a fixed upper bound on the number of juveniles that can be produced by any given parent pair. We will consider a population model where we randomize on *α* in Eq (2). Suppose *ε* ∈ [0, 1), *α*_1_ ∈ (0, 2), and *α*_2_ ≥ 2 are all fixed. In any given generation, the randomly formed parent pairs independently produce juveniles according to Eq (2) with *α* = *α*_1_ with probability *ε*, and with *α* = *α*_2_ with probability 1 − *ε*. Thus, we see the evolution of a highly fecund population evolving according to random sweepstakes as being well described by a mixture of two processes, one which generates only small families (*α* ≥ 2), and another process generating both small and large families and which may then generate large enough families (with 0 < *α* < 2) often enough to affect the evolution of the population (recall Eq (1)). Scalings of *ε* in terms of the population size can be identified so that the model converges (in the appropriate sense) to a non-trivial limit(17). With *X*_1_ (*t*), …, *X*_⌊*M*/2⌋_ (*t*) denoting the random number of juveniles contributed by ⌊*M*/2⌋ parent pairs at time *t*, then *X*_1_ (*t*), …, *X*_⌊*M*/2⌋_ (*t*) are independent and identically distributed, and *X*_1_ (*t*), …, *X*_⌊*M*/2⌋_ (*t*) are independent across generations. Our formulation in Eq (2) guarantees that ℙ (*X* = *j*) ≤ ℙ (*X* = *k*) for 2 ≤ *k* < *j* ≤ *u*(*N*) and ℙ (2 ≤ *X* ≤ *u*(*N*)) = 1.

The incorporation of an upper bound (*u*(*N*) in Eq (2)) on the number of juveniles that can be produced by any given parent pair is biologically reasonable. In a diploid population the juveniles represent (at least) fertilized eggs, not gametes. Even though some organisms, in particular broadcast spawners, are highly fecund, they cannot produce arbitrarily many juveniles. It is highly plausible that for at least some broadcast spawners the number of fertilized eggs produced even over the lifetime of a given individual may be at most a fraction of the population size. For comparison, the capacity to produce arbitrarily large numbers (at least an order of magnitude larger than the population size) of juveniles is necessary to obtain the (complete) Beta(2 − *α*, *α*)-coalescent (a particular example of a multiple-merger coalescent) from the Schweinsberg model(63). Incorporating an upper bound (cutoff) of the form *u*(*N*) = *K N* for some constant *K* > 0 on the number of juveniles each individual can produce can be shown to lead to an incomplete Beta(2 − *α*, *α*)-coalescent(16). Randomizing on *α* solves a timescaling issue with the Schweinsberg model, i.e. the problem of recovering the observed amount of genetic diversity given estimates of *α* without requiring unrealistic assumptions of the population size or mutation rate(17; 24; 26). A further advantage of the population model described here with production of juveniles according to Eq (2) is that it does not only model random sweepstakes, but is also a natural, realistic, and mathematically tractable alternative to the Wright-Fisher model for the case *α* ≥ 2 or *u*(*N*)/*N* → 0 as *N* → ∞(16).

We incorporate recurrent bottlenecks as follows. In any given generation a bottleneck occurs with a fixed probability denoted *b*. If a bottleneck occurs the population is reduced to a fixed number *B* by sampling uniformly at random and without replacement *B* individuals from the current set of diploid individuals. The surviving *B* individuals then produce juveniles as previously described. The law in Eq (2) thus dictates how quickly the population recovers from a bottleneck. The rate of growth after a ‘strong’ (significant reduction in population size) bottleneck has previously been suggested to have to be low for the bottleneck to have a significant effect on genetic variation(56). However, models of strong bottlenecks with a swift recovery have been shown, using coalescent theory to approximate gene genealogies, to lead to significant changes in predictions about genetic diversity(12; 23).

Regardless of a bottleneck or not, if at any time the total number of juveniles does not exceed the carrying capacity *C* they all survive, otherwise we assign a weight to each juvenile and sample *C* juveniles without replacement and with weights. Thus, selection only impacts the evolution of the population when the total number of juveniles exceeds the carrying capacity. Define 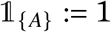 if *A* holds, and zero otherwise. For any time *t* the population size *N*_*t*+1_ is then given by

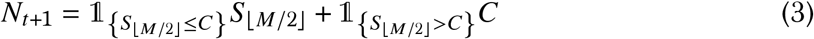

where *S*_⌊*M*/2⌋_ is the total number of juveniles produced by ⌊*M*/2⌋ parent pairs, and

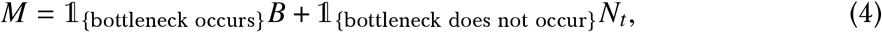

i.e. *M* is the number (*B*) of individuals surviving a bottleneck if a bottleneck occurred, otherwise the number (*N*_*t*_) of individuals from the previous generation. A somewhat similar model of viability selection for a single site in a diploid population evolving according to the Wright-Fisher model but without bottlenecks is considered in Chapter 10 in (28).

We assume the sites are unlinked. Each juvenile inherits the genotypes of its parents according to Mendel’s laws (see § L), we exclude mutation. Each site holds one of two types, either the ‘wild’ type (the type present in the population before the mutation occurs), and the type conferring selective advantage, which we will refer to as the ‘fit’ type. However, since we are modeling a diploid population, the dominance mechanism at each site determines the effect of the fit type on the trait value. A ‘genotype’ is the type obtained by counting the copies of the fit type at a given site. Thus, at each site there are three possible genotypes (*r* (*r* + 1)/2 if *r* types), and 3^*L*^ possible L-site types given *L* sites. We model viability selection by assigning ‘weights’ to the juveniles based on their genotypes. When the number of juveniles exceeds the carrying capacity we sample *C* juveniles without replacement and with weights. Given an L-site type *g* ≔ (*g*_1_, …, *g*_*L*_), where *g*_*ℓ*_ is the genotype at site *ℓ*, assign a weight *w* according to

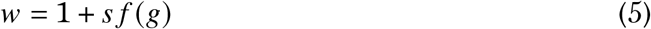

when the effects of the sites are additive, where *s* > 0 is the strength of selection. In§ B we take the weight as

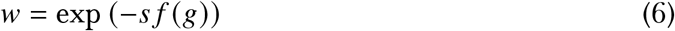

assuming a Gaussian fitness (as for example in (19)) with trait optimum at zero. We refer to the function *f* in Eq (6) and (5) as the weight function. Through the weight function we determine how the genotypes affect the phenotype and set the dominance mechanism at each site. For example, the type conferring selective advantage (the fit type) can be dominant at one site, and semidominant or recessive at other sites. When the number of juveniles exceeds the carrying capacity we sample *C* juveniles by drawing a random exponential with rate the given viability weight (*w*) for each juvenile, and recording the *C*th smallest value *E*_(*C*)_ among all the exponentials; the *C* juveniles with an exponential not exceeding *E*_(*C*)_ survive to form the next generation of reproducing individuals. The relevant stochastic process would be one tracking the number of diploid individuals homozygous for the fit type at all sites. However, one needs to track the number of individuals of all possible *L*-site types since for any site any given pair of parent genotypes can produce from one to all three possible genotypes at the given site.

## 3 Results

We consider various dominance mechanisms (see Table S7 in § J) of the fit type, ranging from overdominance(50), i.e. when the heterozygote is fittest, to the fit type being recessive. The initial configuration is always with the fit type in exactly one copy at each site, i.e. we start with *C* − *L* diploid individuals homozygous for the wild type at all *L* sites, and *L* distinct individuals each heterozygous for the fit type at a distinct site, and homozygous for the wild type at all other sites.

We define ‘fixation’ to have occurred when the fit type has fixed at all sites. We therefore need to track the genotypes at all sites, since if a fit type is lost at one site fixation by definition cannot occur. The time to fixation, conditional on fixation occurring, is taken to be the first time fixation happens. Let {*Y*_*t*_ } taking values in 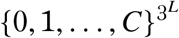 denote the process tracking the number of diploid individuals of each of the *L*-site types *g* ∈ *G*, with *G* denoting the set of all possible L-site types. Let {*Z*_*t*_ } taking values in [0, 1]^*L*^ denote the process tracking the frequency of the fit type at each site. The values of {*Z*_*t*_ } can be obtained from {*Y*_*t*_ } by summing over the L-site types. With *Y*_*t*,*g*_ denoting the number of individuals of *L*-site type *g* at time *t*

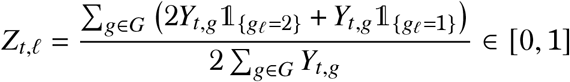

is the relative frequency of the fit type at site *ℓ* at time *t*. Let, for *x* ∈ [0, 1],

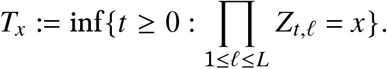

Then the time when the fit type has been lost for at least one site is *T*_0_, and *T*_1_ the time when the fit type has fixed at all sites, with the understanding that we are starting with one copy of the fit type at each site as already described. We can then define the probability of fixation

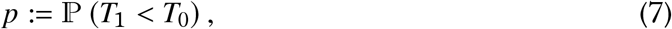

In the presence of recurrent bottlenecks a type may become fixed (or lost) in the population at any population size *N*_*t*_ ∈ {*B*, *B* + 1, …, *C*}. We are also interested in the time to fixation conditional on fixation with expected value

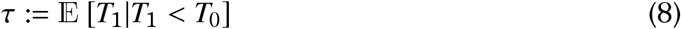

We assess the effects of recurrent bottlenecks, random sweepstakes, varying dominance mechanisms and epistasis on selection through the effects on the fixation trajectories and on the statistics *p* and *τ*. The statistics *p* and *τ* are key statistics for evaluating the impact of selection on a population. We do not aim for precise estimates of *p* or *τ*, rather to identify general ways in how the stated elements may affect selection when acting at more than one site. We report the results in the form of fixation trajectories shown as number of diploid individuals homozygous for the fit type at all sites in question each time relative to the population size. Graphs in right columns (e.g. Figure 1; Figure S6 in § B.1) plot each excursion from the corresponding graph on the left with the time of each excursion normalised by the time to fixation for the excursion. Corresponding numerical estimates of *p* and *τ* are recorded in § D. When comparing the graphs within each figure note that the scale of the abscissa (time axis) may vary between graphs. The algorithm is given in the form of a pseudo-code in § K.

**Figure 1:**
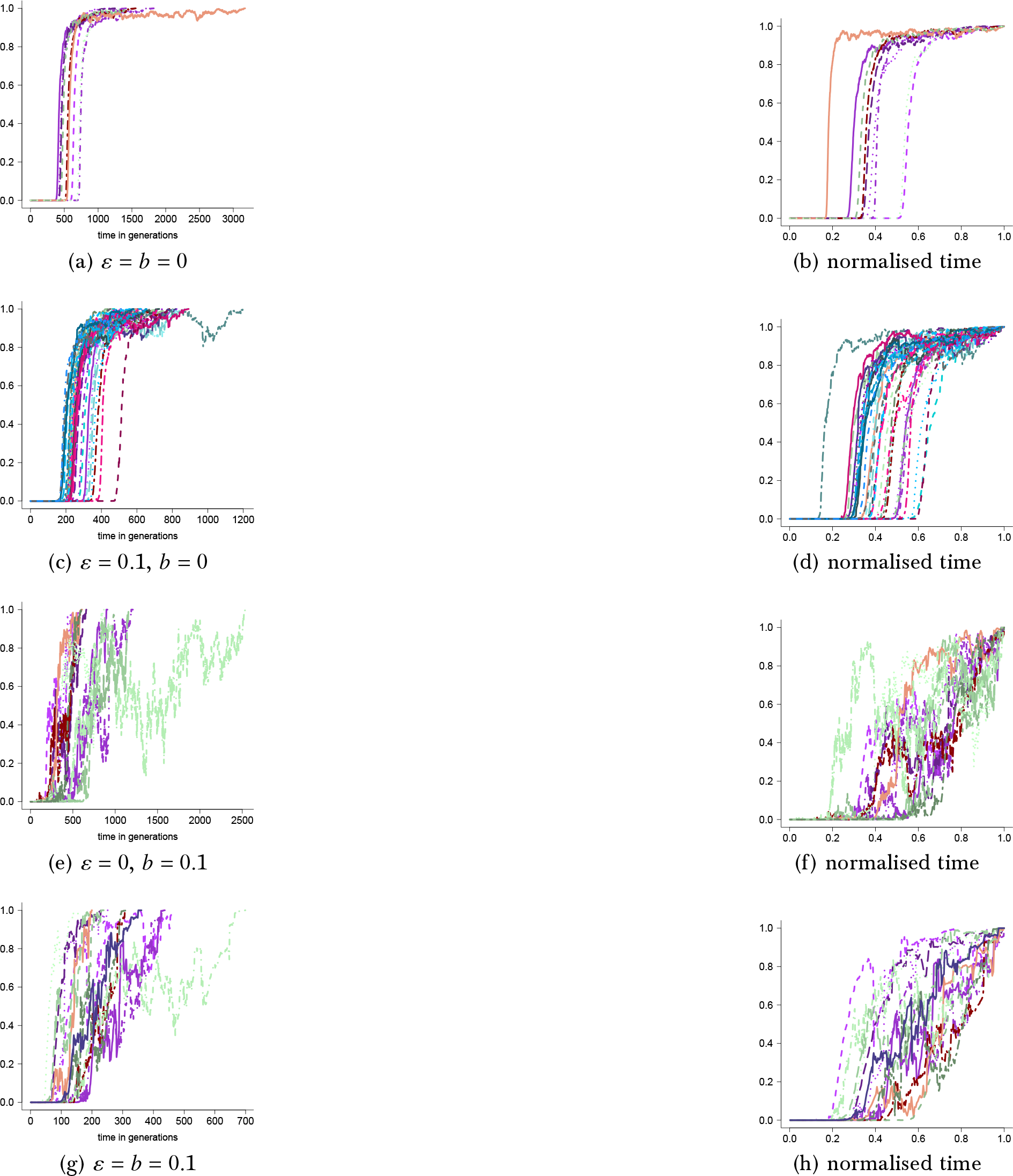
Examples of excursions to fixation of the fit type at two sites shown as number of diploid individuals homozygous for the fit type at both sites relatives to the population size as a function of time with weight computed as in Eq (5) with weight function as in Eq (9) for carrying capacity *C* = 10^4^ and *α*_1_ = 0.75, *α*_2_ = 3 with selection strength *s* = 1 and *B* = 10^2^ and *ε* and *b* as shown. The panels on the right show the excursions from the left side panels with the time for each excursion normalised by the time to fixation for the excursion. The corresponding numerical estimates of *p* and *τ* are recorded in Table S1. The scale of the abscissa (time axis) may vary between the panels

### 3.1 Two sites

Even though our algorithm works for any number of unlinked sites, a natural starting point is to investigate the effects of random sweepstakes and bottlenecks on selection for two sites (*L* = 2). For each site, genotype 0 (0/0) is homozygote for the wild type, genotype 1 (0/1) is the heterozygote, and genotype 2 (1/1) is homozygote for the fit type. We write

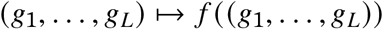

to denote how the positive bounded weight function, denoted *f* in Eq (5) (and Eq (6)), is computed as a function of a given *L*-site type (*g*_1_, …, *g*_*L*_) ∈ {0, 1, 2}^*L*^.

With the weight computed as in Eq (5) consider the weight function given by

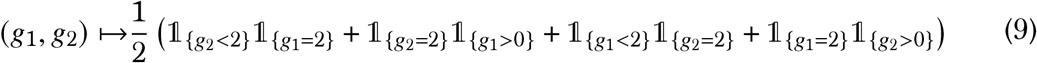

The weight function in Eq (9) is an example of a symmetric epistasis where the fit type at site *ℓ*_1_ is recessive 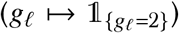 if there is at most one copy of the fit type at site *ℓ*_2_, otherwise the fit type is dominant 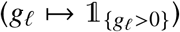 at site *ℓ*_1_. See Figure 1 for examples of fixation trajectories, where the panels on the right show the excursions with the time of each excursion normalised by the time to fixation for the excursion. In Figure S7 in § B.2 we give another brief example for a similar form of epistasis for carrying capacity *C* = 10^6^.

The effect of epistasis as defined by Eq (9) can be assessed by comparing Figure 1 to Figure 2, Figure 3, and Figure 4 (the corresponding numerical estimate of *p* and *τ* are recorded in Table S1). The results in Figure 2 (see Figure S1 for a comparison to Figure 2a and Fig 2b with carrying capacity *C* = 10^5^) hold for the weight computed as in Eq (5) with the weight function given by

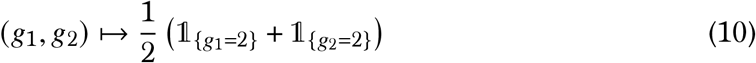

**Figure 2:**
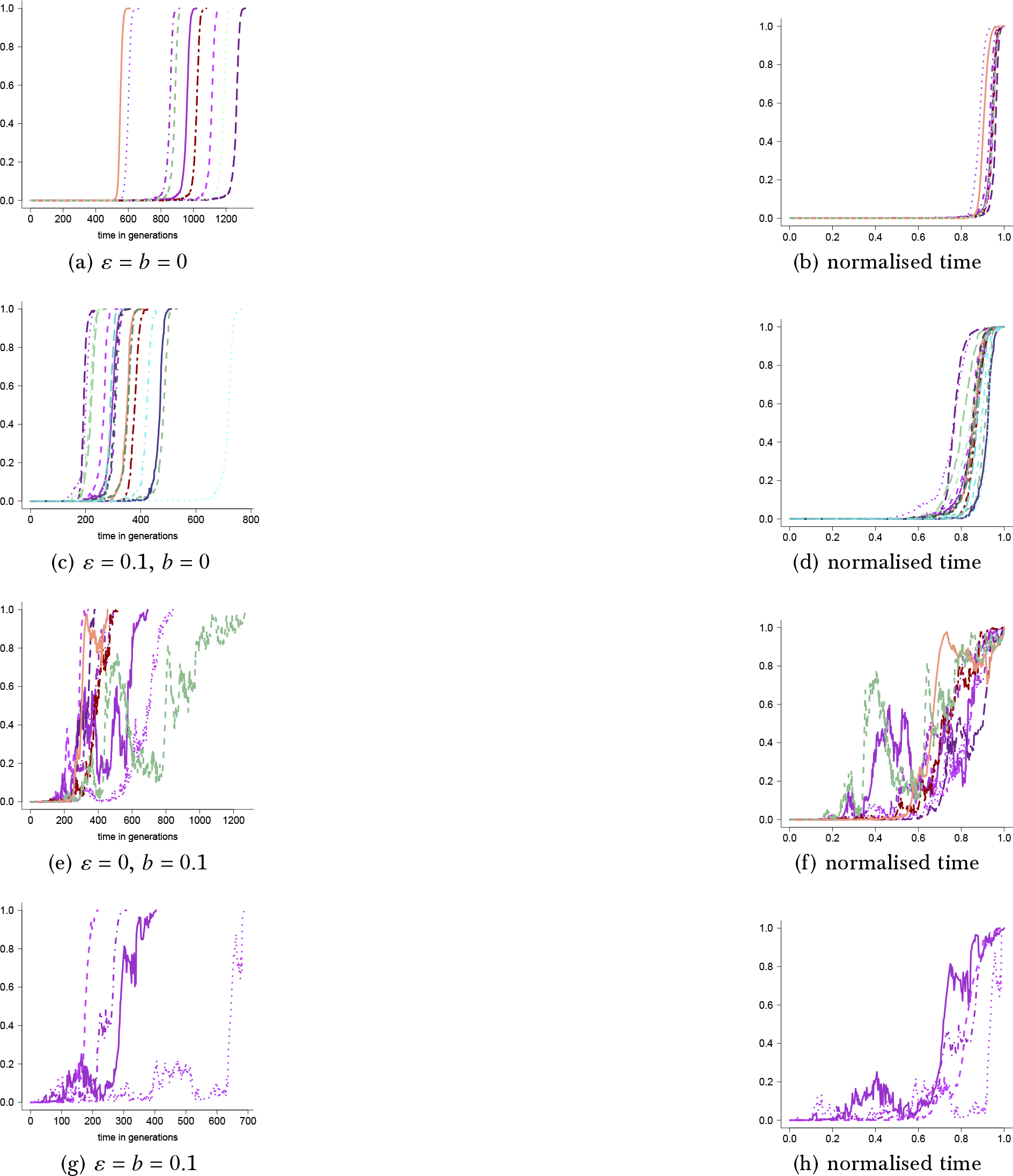
Examples of excursions to fixation for the fit type at two sites shown as the number of diploid individuals homozygous for the fit type at both sites relative to the population size as a function of time with weight computed as in Eq (5) with weight function as in Eq (10) for carrying capacity *C* = 10^4^, strength of selection (Eq (5)) *s* = 1, *α*_1_ = 0.75, *α*_2_ = 3 with *ε* and *b* as shown and bottleneck size *B* = 10^2^. Corresponding estimates of *p* and *τ* are recorded in Table S1; the scale of the abscissa (time axis) may vary between the panels

**Figure 3:**
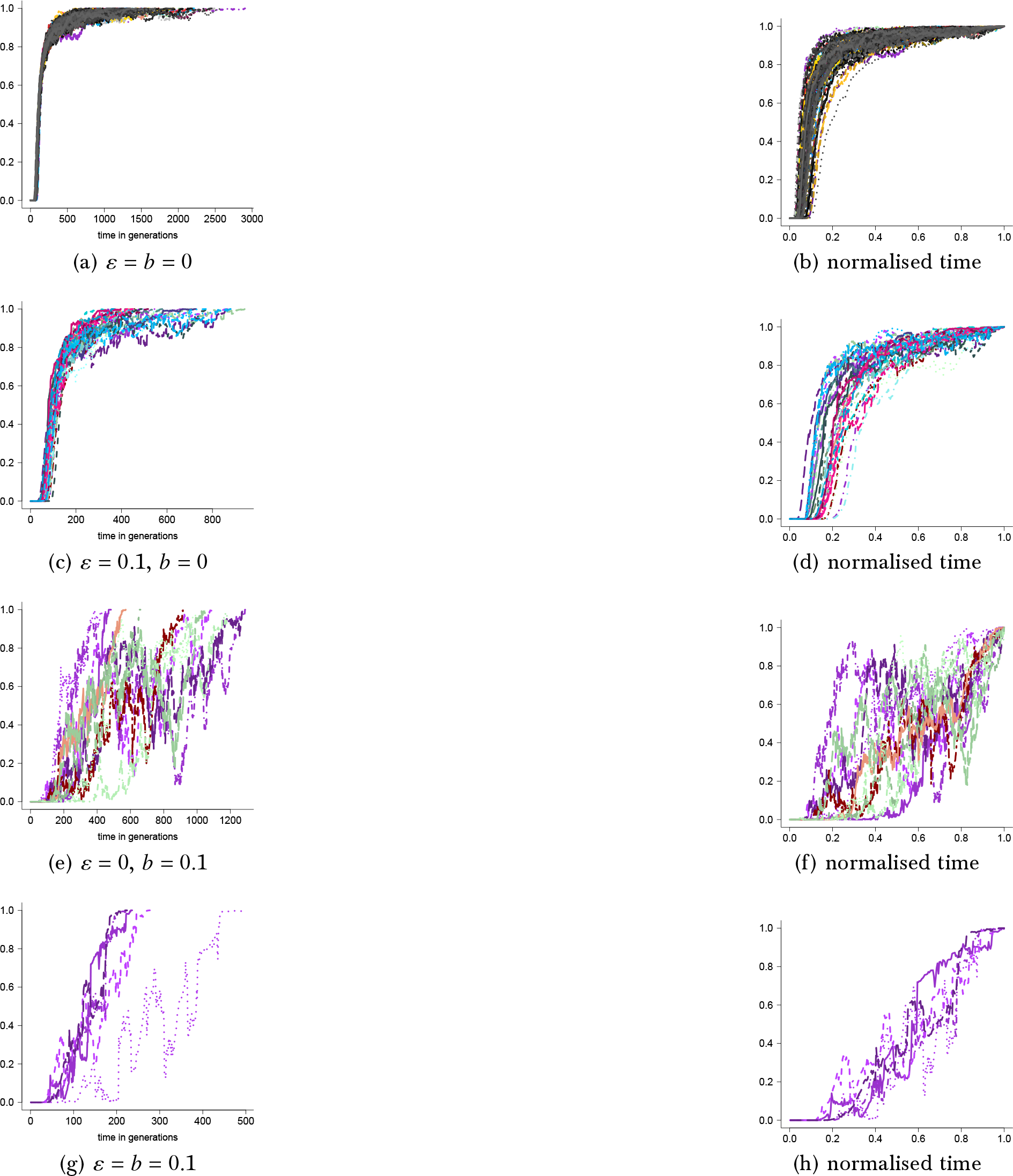
Examples of excursion to fixation shown as the number of diploid individuals homozygous for the fit type at both sites as a function of time relative to the population size for the fit type dominant at both sites with weight computed as in Eq (5) with weight function as in Eq (11) for carrying capacity *C* = 10^4^, *α*_1_ = 0.75, *α*_2_ = 3, cutoff *u*(*N*) = *C*, strength of selection (Eq (5)) *s* = 1, bottleneck size *B* = 10^2^ with *ε* and *b* as shown. Corresponding estimates of *p* and *τ* are recorded in Table S1. The scale of the abscissa (time axis) may vary between the graphs

**Figure 4:**
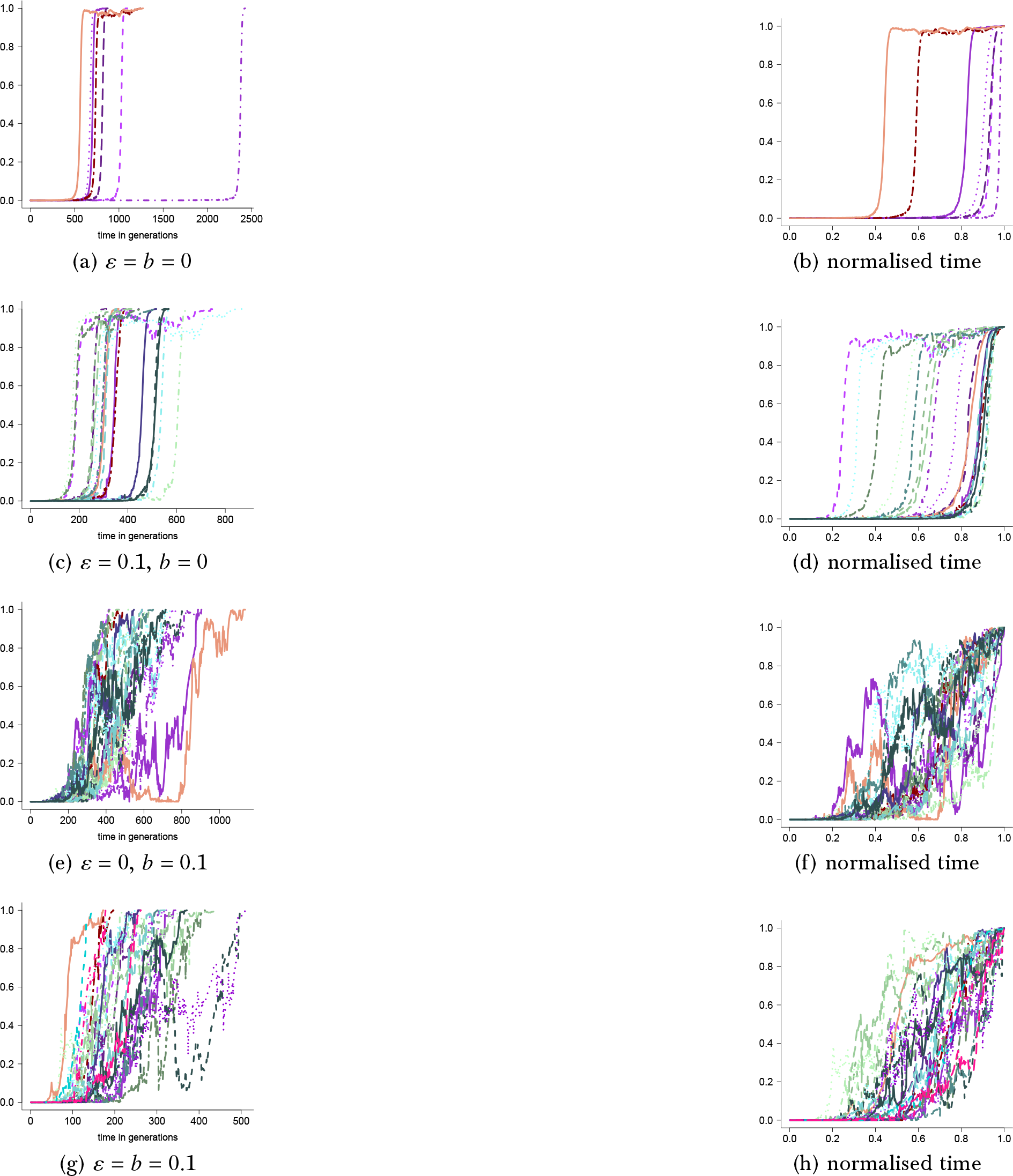
Examples of excursions to fixation for two sites shown as number of diploid individuals homozygous for the fit type at both sites relative to the population size as a function of time for weight computed as in Eq (5) with weight function given as in Eq (12) for carrying capacity *C* = 10^4^, *α*_1_ = 0.75, *α*_2_ = 3, strength of selection (Eq (5)) *s* = 1, bottleneck size *B* = 10^2^, *ε* and *b* as shown. In the panels on the right the time of each excursion is normalised by the time to fixation for the excursion. Corresponding estimates of *p* and *τ* are recorded in Table S1. The scale of the abscissa (time axis) may vary between the panels

**Figure 5:**
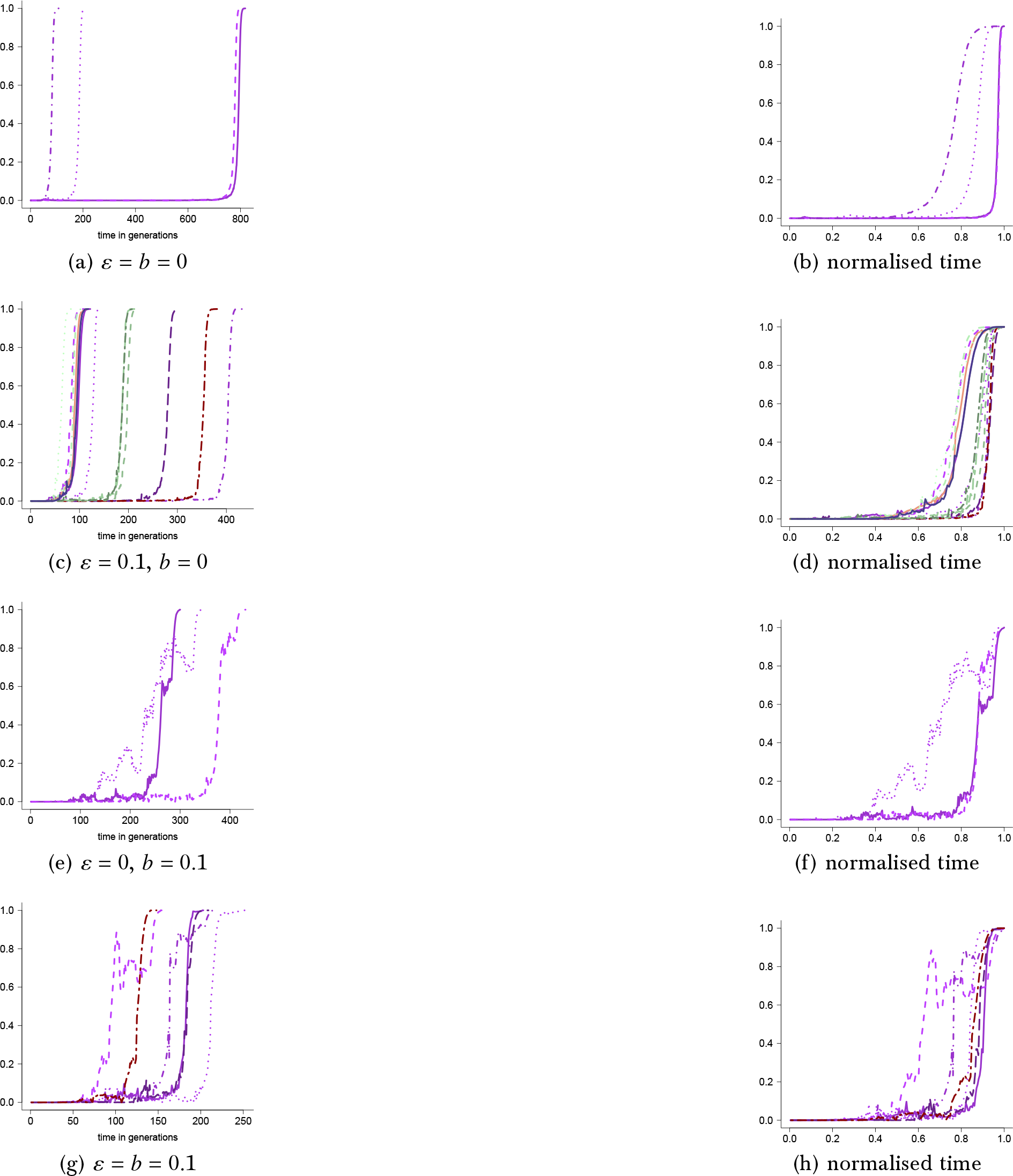
Examples of excursions to fixation for two sites shown as the number of diploid individuals homozygous for the fit type at both sites relative to the population size as a function of time with multiplicative fitness with weight computed as in Eq (14) for carrying capacity *C* = 10^4^, cutoff *u*(*N*) = *C*, *α*_1_ = 0.75, *α*_2_ = 3, *B* = 10^2^ and *ε* and *b* as shown. The time of each excursion in the right panels is normalised by the time to fixation for the excursion. Corresponding estimates of *p* and *τ* are in Table S2. The scale of the abscissa (time axis) may vary between graphs

**Figure 6:**
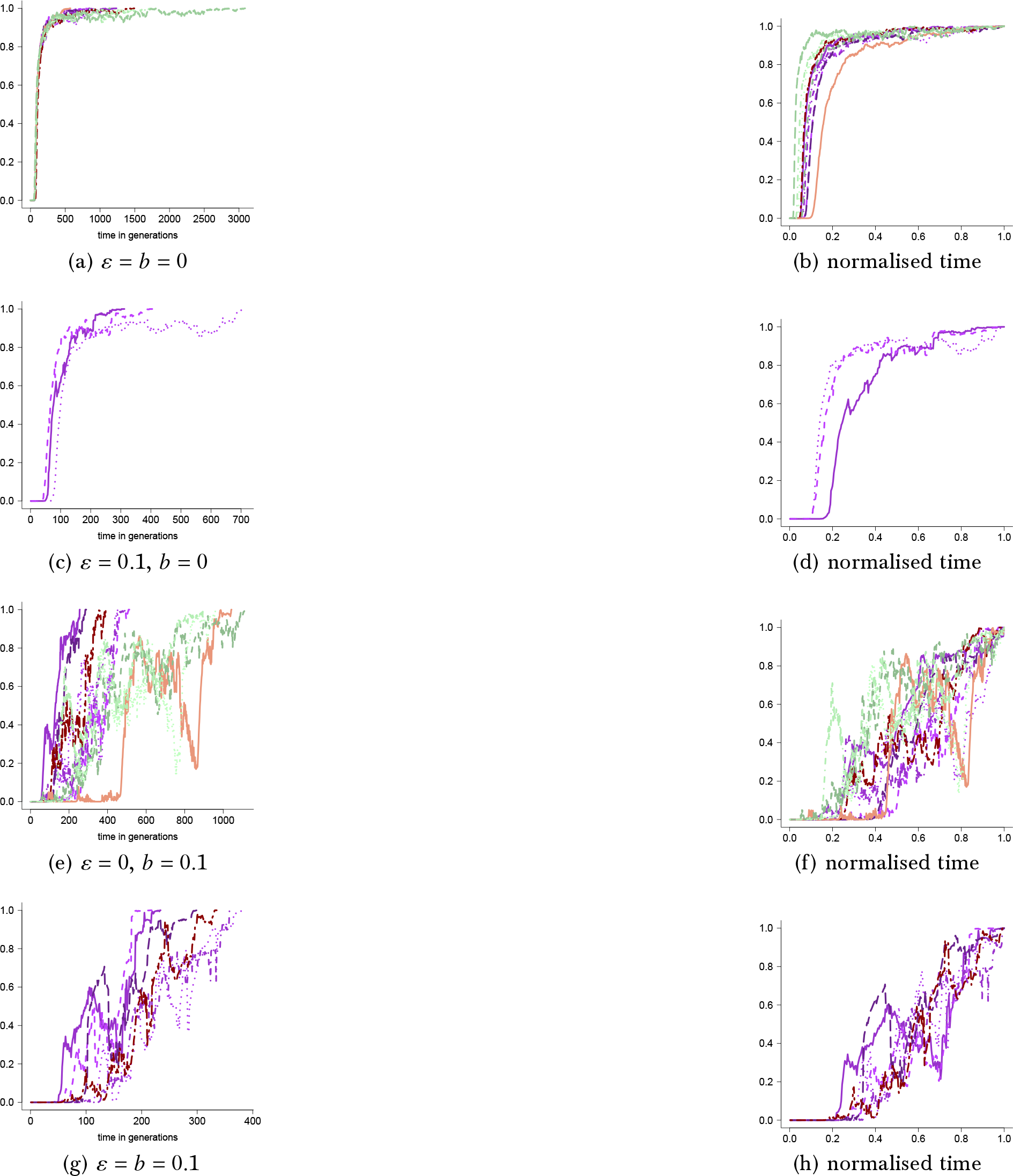
Examples of excursions to fixation for two sites shown as the number of diploid individuals homozygous for the fit type at both sites relative to the population size as a function of time with multiplicative fitness with weight computed as in Eq (15) for carrying capacity *C* = 10^4^, cutoff *u*(*N*) = *C*, *α*_1_ = 0.75, *α*_2_ = 3, cutoff *u*(*N*) = *C*, strength of selection *s* = 1, bottleneck size *B* = 10^2^ and *ε* and *b* as shown. The time of each excursion in the right panels is normalised by the time to fixation for the excursion. Corresponding estimates of *p* and *τ* are recorded in Table S2. The scale of the abscissa (time axis) may vary between graphs

**Figure 7:**
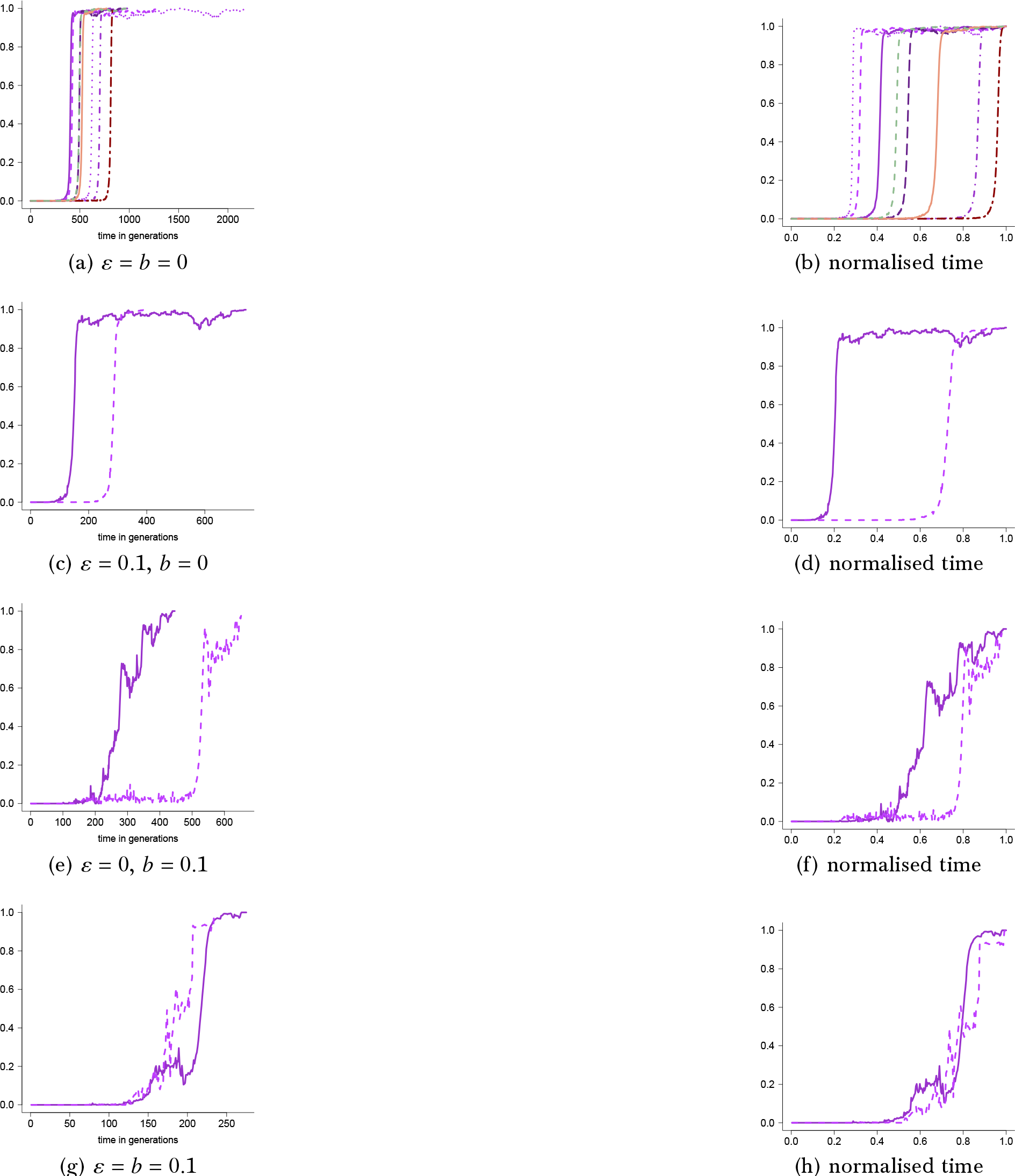
Examples of excursions to fixation for two sites shown as the number of diploid individuals homozygous for the fit type at both sites relative to the population size with multiplicative fitness with weight computed as in Eq (16) with carrying capacity *C* = 10^4^, *α*_1_ = 0.75, *α*_2_ = 3, strength of selection *s* = 1, bottleneck size *B* = 10^2^ and *ε* and *b* as shown. The time of each excursion in the right panels is normalised by the time to fixation for the excursion. Corresponding numerical estimates of *p* and *τ* are recorded in Table S2. The scale of the abscissa (time axis) may vary between graphs

**Figure 8:**
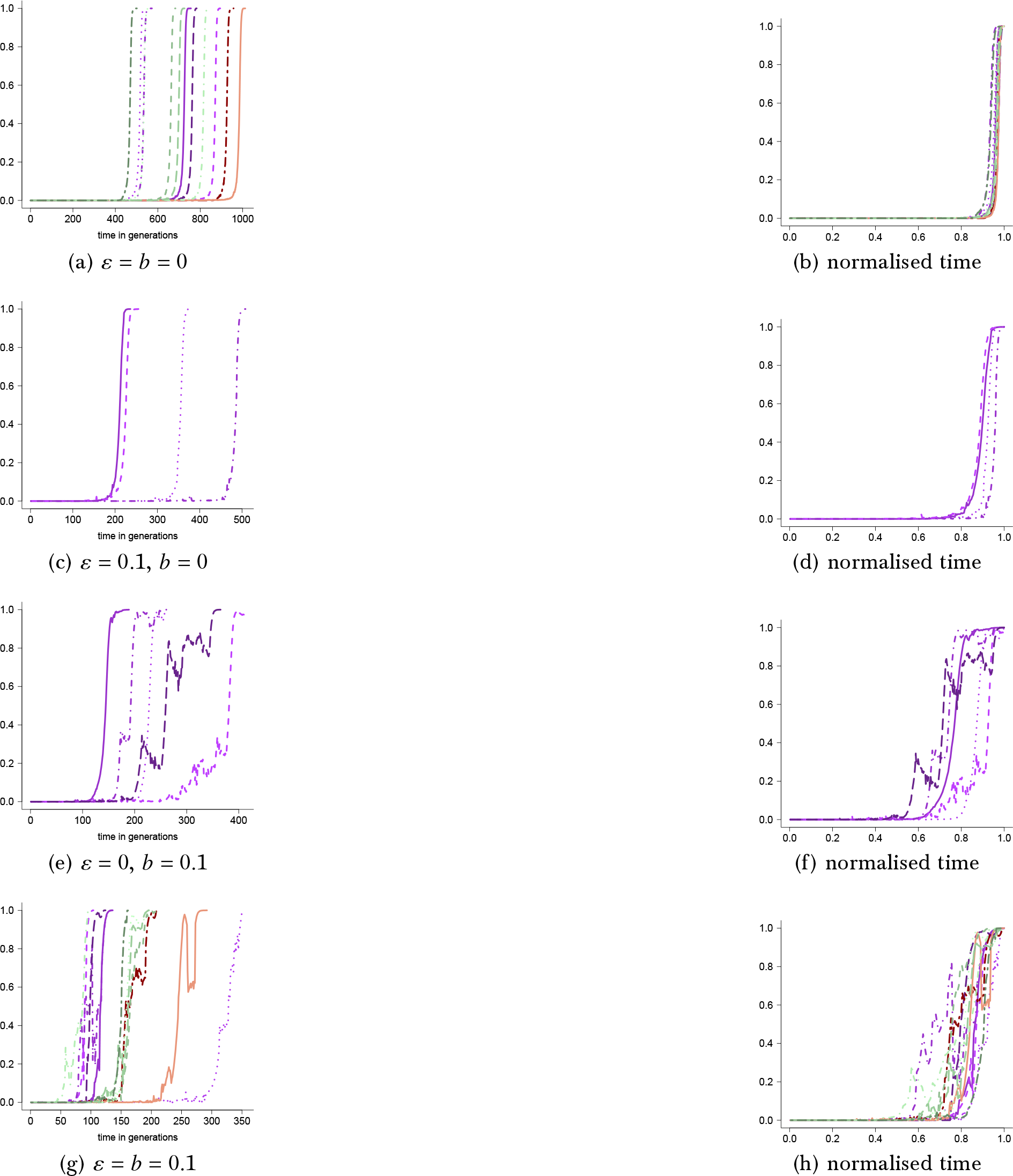
Examples of excursions to fixation at two sites shown as number of diploid individuals homozygous for the fit type at both sites relative to the population size as a function of time for weight as in Eq (17) with carrying capacity *C* = 10^4^, strength of selection *s* = 1, *α*_1_ = 0.75, *α*_2_ = 3, cutoff *u*(*N*) = *C*, bottleneck size *B* = 10^2^, *ε* and *b* as shown. The time of each excursion in the right panels is normalized by the time to fixation for the excursion. The corresponding estimates of *p* and *τ* are in Table S2. The scale of the abscissa (time axis) may vary between panels

Eq (10) states that the fit type is recessive at both sites and there is no epistasis. The results in Figure 2 can be compared to Figure S5, in which the weight is given by Eq (6) and the carrying capacity is *C* = 10^6^. In Figure 3 the weight is computed according to Eq (5) with the weight function given by

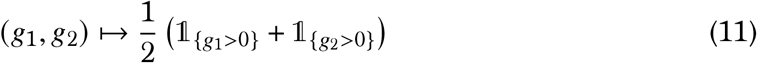

In Eq (11) the fit type is dominant at both sites and there is no epistasis. The results in Figure 3 can be compared to Figure S2 in § B, in which the weight is given by Eq (6) and the carrying capacity is *C* = 10^6^ (see also Figure S3).

Figure 4 holds results for the weight computed as in Eq (5) with the weight function given by

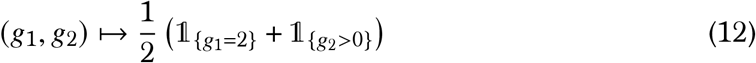

In Eq (12) the fit type is dominant at one site and recessive at the other site without epistasis. The results in Figure 4 can be compared to Figure S4, which holds results for the weight computed as in Eq (6) with carrying capacity *C* = 10^6^.

With epistasis as defined in Eq (9), the recessive mechanism is clearly decisive regarding *p* (see Table S1), while there is seems no significant effect of epistasis on *τ*. Interestingly, the fixation trajectories when both dominance and recessive mechanisms are involved (Fig 1a, Fig 4a) show something of a mixture effect between only recessive (Fig 2a) and only dominance (Fig 3a) mechanisms. Random sweepstakes and bottlenecks reduced both *p* and *τ* relative to the absence of both (Table S1).

In Figure 1–Fig 4 we consider a additive fitness model (e.g. Eq (9)). In Figures 5–Fig 8 we consider multiplicative fitness models (see Table S2 for the corresponding numerical estimates of *p* and *τ*) where the weight is computed according to the general formulation, with *g* ≡ (*g*_1_, …, *g*_*L*_),

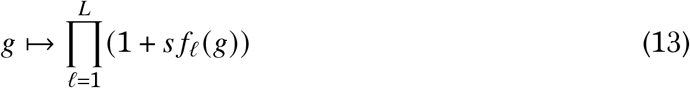

where *s* ≥ 0 is again the strength of selection. The functions *f*_*ℓ*_ (*g*) are projections onto the *ℓ*th coordinate of (*g*_1_, …, *g*_*L*_) taking into account possible epistatic interactions. Furthermore in Figures 5–Fig 8 we restrict to the fit type being either dominant or recessive for comparison with Figure 1–Fig 4. In Fig 5 we consider a simple multiplicative fitness model of symmetric epistasis, where the weight is computed as

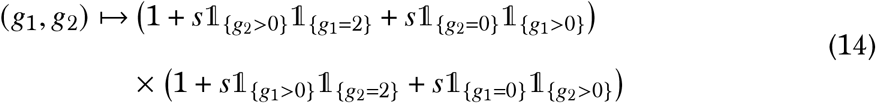

In Eq (14) the fit type is recessive at site *ℓ*_1_ if there is at least one copy of the fit type at site *ℓ*_2_, otherwise the fit type is dominant at site *ℓ*_1_. For comparison we also consider multiplicative fitness without epistasis (Figures 6–Fig 8). In Figure 6 the weight is given by

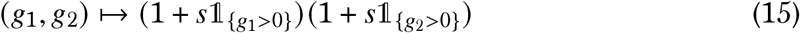

In Eq (15) the fit type is dominant at both sites without epistasis. In Figure 7 with weight as given by

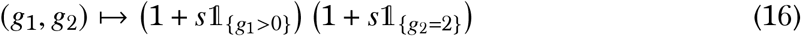

In Eq (16) the fit type is dominant at one site and recessive at the other site. In Figure 8 the weight is given by

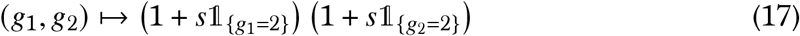

In Eq (17) the fit type is recessive at both sites and there is no epistasis.

In contrast to the additive model (Eq (9)) epistasis in the multiplicative model predicts in most cases shorter time to fixation (conditional on fixation) compared to the models of multiplicative fitness without epistasis (Table S2). Furthermore, also in contrast with the additive model, the probability of fixation is on the higher side in comparison with the models of multiplicative fitness without epistasis. For both the additive and the multiplicative model we see that the fixation trajectories can be mixtures (e.g. Fig 4b, Fig 7b) of the trajectories associated for strictly dominant (Fig 3, Fig 6) and strictly recessive (Fig 2, Fig 8) mechanisms. We also remark that in the absence of strong recurrent bottlenecks the fit type when conditioned to fix spends most of the time to fixation in either very low or very high frequency, and very little time (in comparison) in intermediate frequency. However, including both recurrent bottlenecks and random sweepstakes the type may spend significant time in intermediate frequency (e.g. Fig 3h, Fig 6h). In summary, random sweepstakes and bottlenecks reduce both *p* and *τ* when the fit type is either dominant or recessive; random sweepstakes do not really alter the fixation trajectories, but the recurrent strong bottlenecks clearly do.

We have considered the effect of epistasis on selection at two sites when restricting to the fit type being either dominant or recessive (Figures 1–Fig 8; Table S1, Table S2). Now we turn to results when the fit type at (at least) one of the sites is overdominant (Table S7).

Assuming the fit type is overdominant at both sites we consider additive fitness models where the weight function in Eq (5) is given by

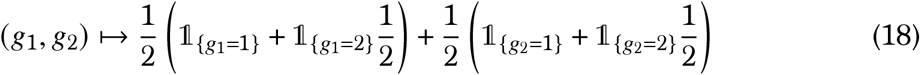

In Eq (18) the fit type is overdominant at both sites without epistasis and each trial resulted either in a loss of the fit type, or an apparent stationary distribution of the frequency of the fit type at both sites (Figure 9; the excursions labelled OO are for the model in Eq (18)). In addition to the fit type being overdominant at both sites as in Eq (18) we also consider mixed cases given by the weight function from Eq (5) as

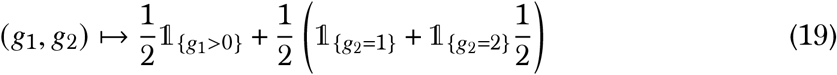

where the fit type absent epistasis is dominant at one site and overdominant at the other site (Figure 9a),

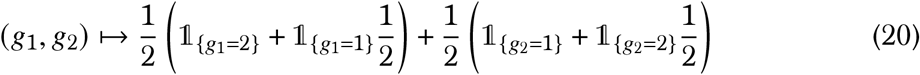

in which the fit type is semidominant at one site and overdominant at the other site without epistasis (Figure 9b), and

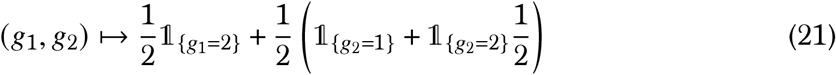

in which the fit type is recessive at one site and overdominant at the other site without epistasis (Figure 9c). The scale of the abscissa in Figure 9 is relative to the carrying capacity. In the absence of random sweepstakes overdominance is known to maintain genetic variability(50). The results in Figure 9 suggest that overdominance can maintain genetic variability also in the presence of random sweepstakes. However, the presence of recurrent bottlenecks only resulted in a loss of the fit type.

**Figure 9:**
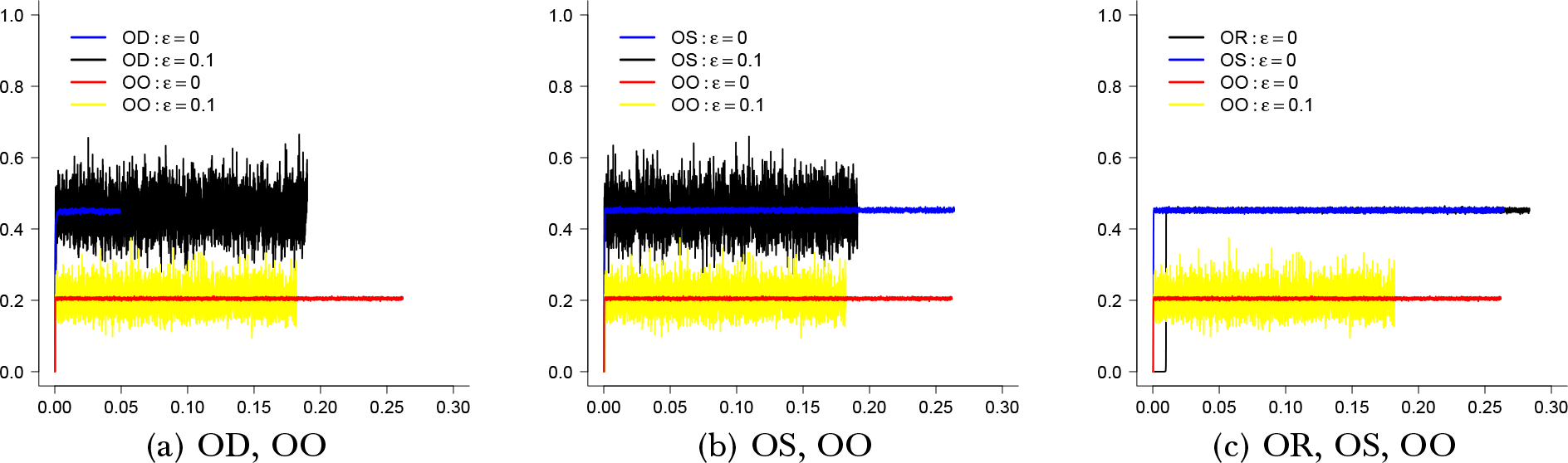
Examples of trajectories for two sites shown as the number of individuals at time homozygous for the fit type at both sites relative to the population size as a function of time with carrying capacity *C* = 10^6^, cutoff *u*(*N*) = *C*, *α*_1_ = 1.05, *α*_2_ = 3, strength of selection *s* = 0.5 and no bottlenecks (*b* = 0) throughout with *ε* as shown and when at least one site the fit type is overdominant with the weight function *f* in Eq (5) given by Eq (18) when the fit type is overdominant at both sites (OO), or the fit type at one site dominant (OD) with the weight function as in Eq (19), semidominant (OS) with the weight function as in Eq (20), recessive (OR) with the weight function as in Eq (21). The scale of the abscissa (time axis) is relative to the carrying capacity

In light of the results related to Figure 9, one might ask if mixtures of over- and underdominance might lead to fixation (granted that such mixtures are realistic). Consider a random additive fitness landscape model of two sites involving overdominance and epistasis, where the weight function in Eq (5) is given by (compare with Eq (33) in § C where the randomness only holds for one site)

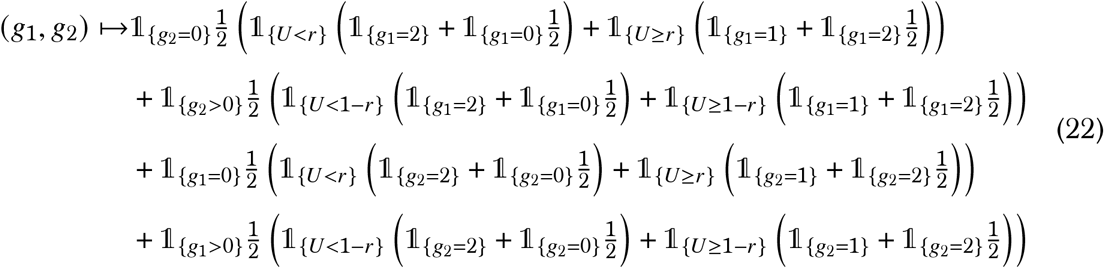

where *r* ∈ (0, 1) is a constant and *U* a random uniform on the unit interval. A diagram illustrating the mechanism in Eq (22) is shown in Eq (23), where 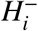 denotes genotype 0/0 (homozygous for the wild type) at site *i*, and 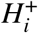 denotes the presence of at least one copy of the fit type at site *i*.

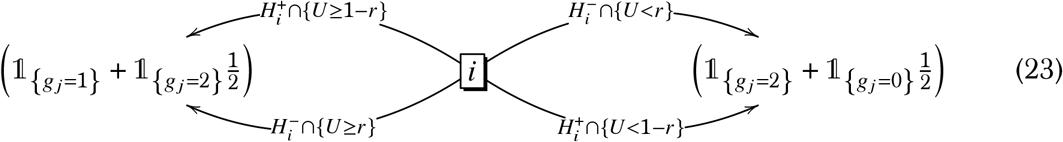

In the the symmetric epistasis modeled in Eq (22), the fit type is underdominant 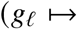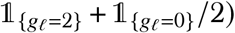 at one site with probability *r*, otherwise overdominant 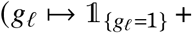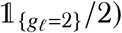, if the type is absent from the other site. On the other hand, the fit type is underdominant with probability 1 − *r*, otherwise overdominant, at one site in case there is at least one copy of the type at the other site. In contrast to the cases without epistasis (Fig 9), given the model in Eq (22), one obtains trajectories to fixation (Figure 10; the corresponding estimates of *p* (Eq (7)) and *τ* (Eq (8)) are recorded in Table S3). Trajectories to fixation are also obtained for a similar model (Eq (33); see Figure S9 and corresponding numerical estimates of *p* and *τ* in Table S6).

**Figure 10:**
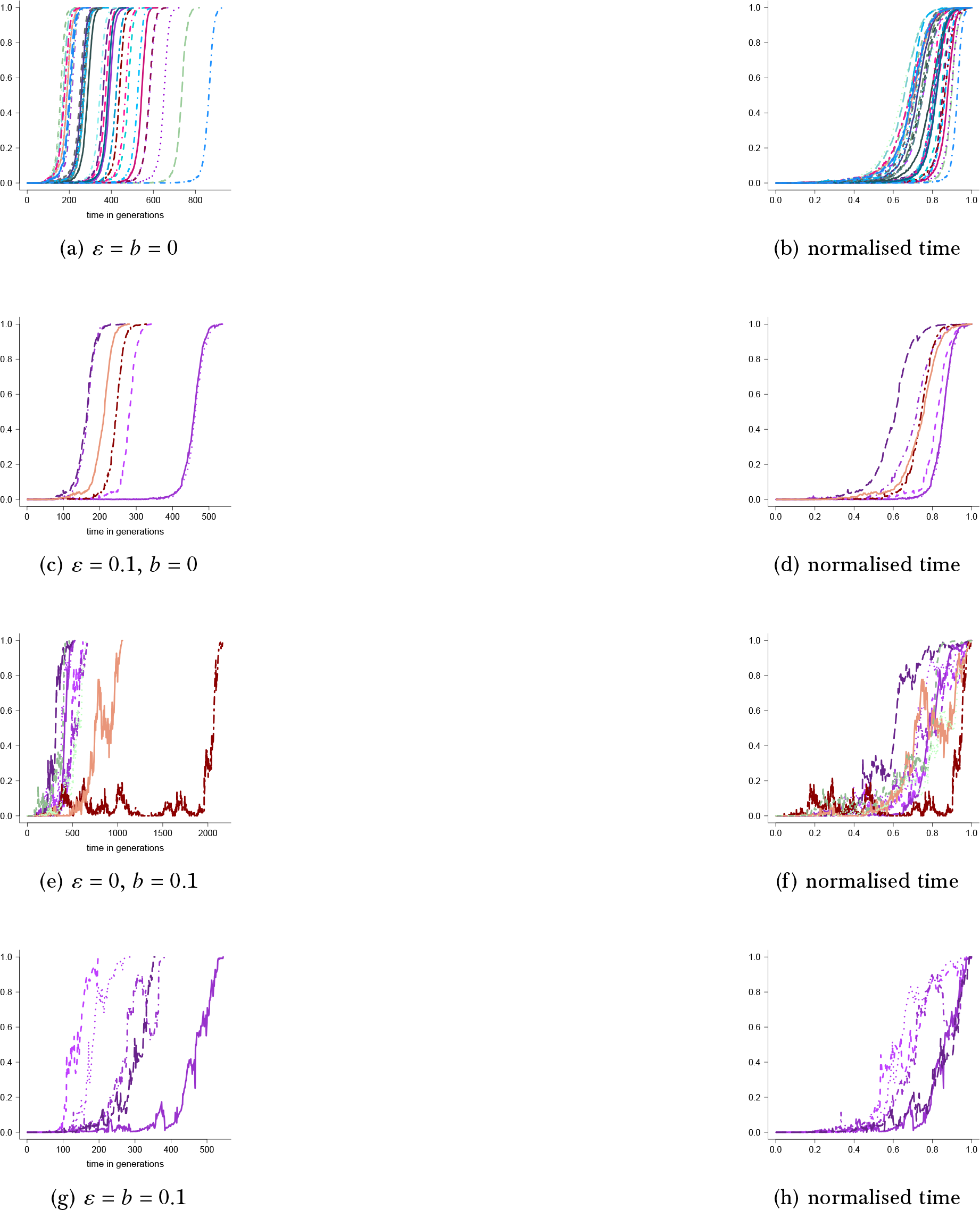
Examples of excursion to fixation at two sites shown as number of diploid individuals homozygous for the fit type at both sites relative to the population size as a function of time when there is epistasis among the sites as in Eq (22), see diagram Eq (23), with carrying capacity *C* = 10^4^, *α*_1_ = 0.75, *α*_2_ = 3, cutoff *u*(*N*) = *C*, and strength of selection (Eq (5)) *s* = 1, bottleneck size *B* = 10^2^, and *ε* and probability of a bottleneck *b* as shown. The time of each excursion in the right panels is normalized by the time to fixation for the excursion. The corresponding numerical estimates of *p* and *τ* are in Table S3. The scale of the abscissa (time axis) may vary between panels

For comparison with the additive fitness model in Eq (22) we consider a multiplicative fitness landscape (recall Eq (13)) restricted to under- and overdominance of the fit type where *r* ∈ (0, 1) is a constant, *U* a random uniform on the unit interval, and

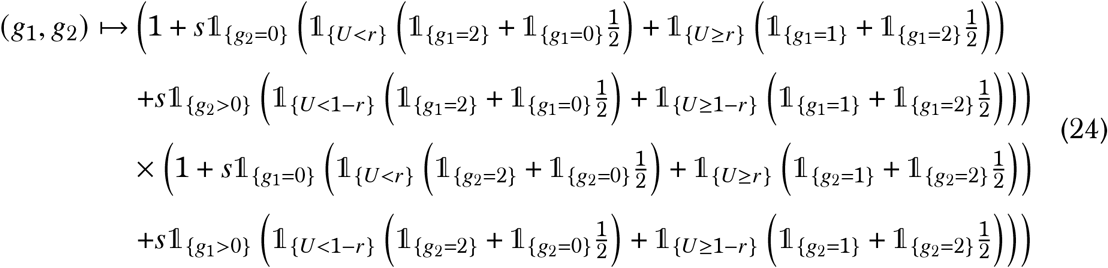

The fitness landscape model in Eq (24) is simply the one in Eq (22) adapted to multiplicative fitness (recall Eq (13)). As for the additive model, the multiplicative model (Eq (24)) also yields excursions to fixation (Figure 11). In Figure 12 the weight is computed as in Eq (24) with the exception that the probability of overdominance or underdominance at a given site depends only on the configuration ({*g*_*ℓ*_ = 0} vs. {*g*_*ℓ*_ > 0}) at the same site, and is independent of the configuration at the other site. In summary, for the models involving under- and overdominance, random sweepstakes reduce both *p* and *τ*, although the effects are quite moderate. In contrast to the models where the fit type was either dominant or recessive, bottlenecks increase *τ* (Table S3).

**Figure 11:**
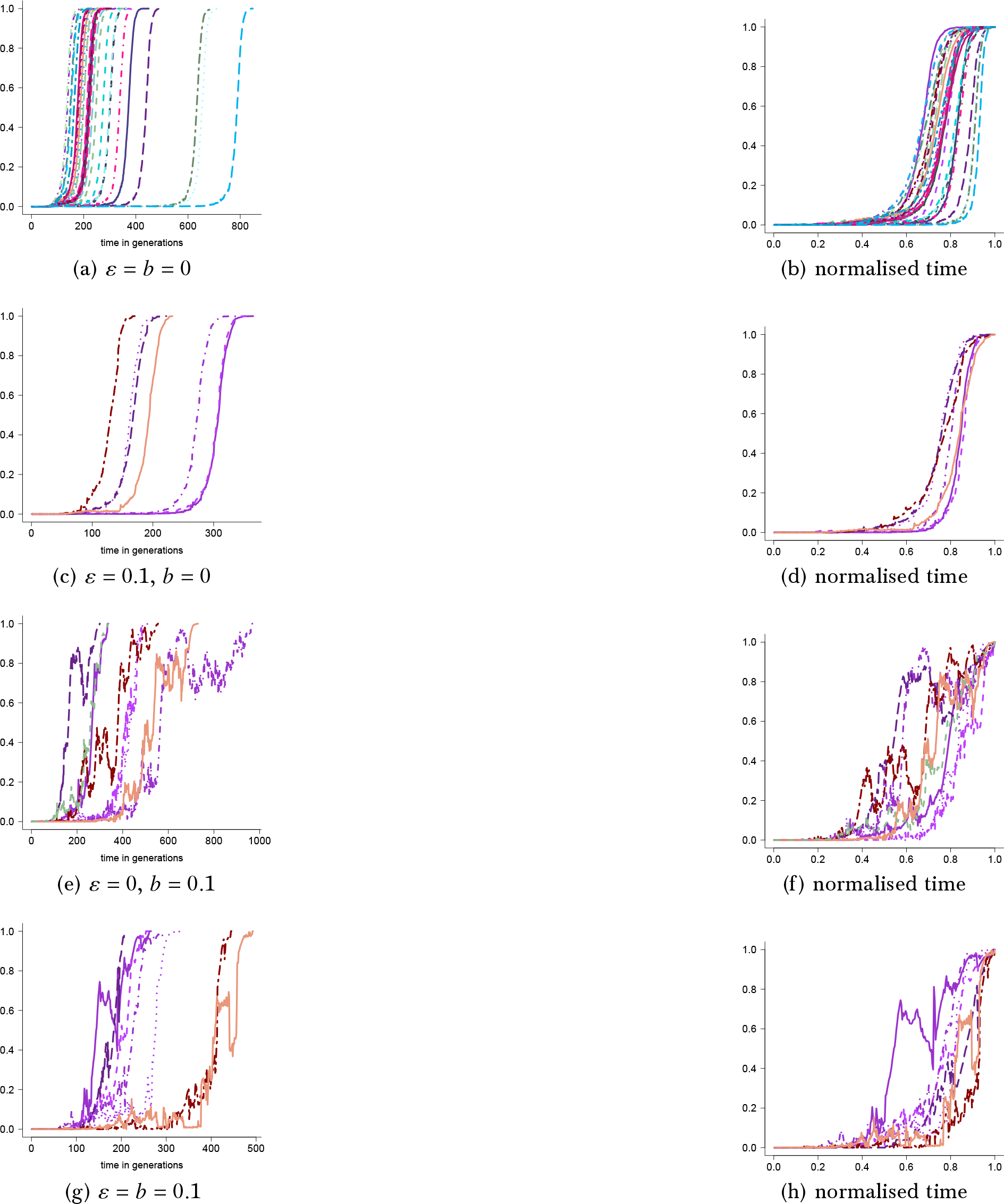
Examples of excursion to fixation for two sites shown as number of diploid individuals homozygous for the fit type at both sites relative to the population size as a function of time when there is epistasis among the sites as in Eq (24) with *r* = 0.25, carrying capacity *C* = 10^4^, *α*_1_ = 0.75, *α*_2_ = 3 and strength of selection *s* = 1 with bottleneck size *B* = 10^2^ and *ε* and probability of a bottleneck *b* as shown. The time of each excursion in the right panels is normalized by the time to fixation for the excursion. Corresponding estimates of *p* and *τ* are recorded in Table S3. The scale of the abscissa (time axis) may vary between panels

**Figure 12:**
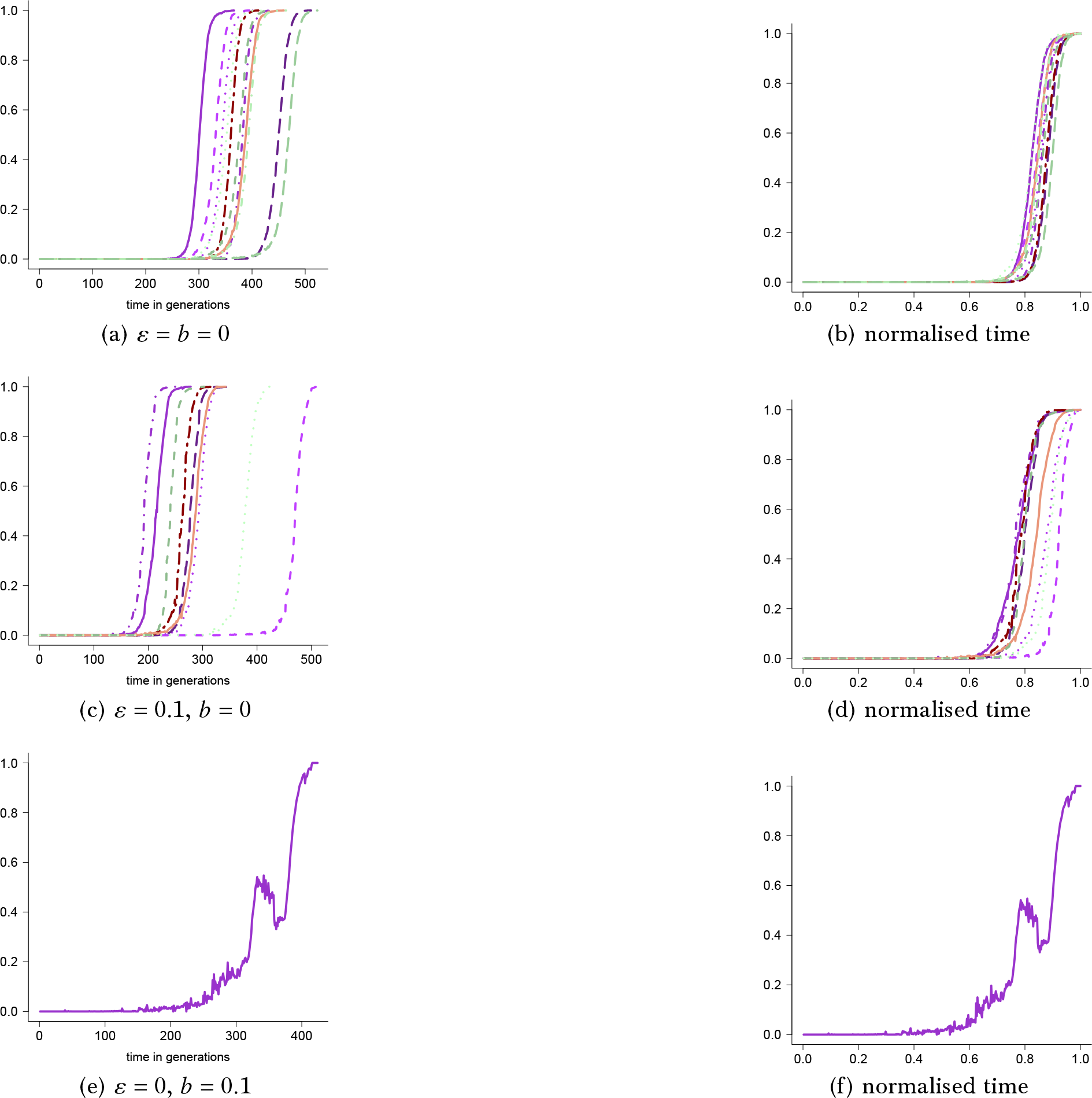
Examples of excursion to fixation for two sites shown as number of diploid individuals homozygous for the fit type at both sites relative to the population size when the weight is computed as in Eq (24) but without epistasis, i.e. the probability of underdominance or overdominance at either site depends only on the configuration (*g*_*ℓ*_ = 0 vs. *g*_*ℓ*_ > 0) at the same site, not on the configuration at the other site; carrying capacity *C* = 10^4^, *α*_1_ = 0.75, *α*_2_ = 3 and strength of selection *s* = 1 with bottleneck size *B* = 10^2^ and *ε* and probability of a bottleneck *b* as shown. The time of each excursion in the right panels is normalized by the time to fixation for the excursion.

### 3.2 Four sites

It is clearly of interest to investigate if fixation at more than two sites can occur (see e.g. Fig S8 in § C). To this end we extend the additive model in Eq (22) to *L* sites to obtain

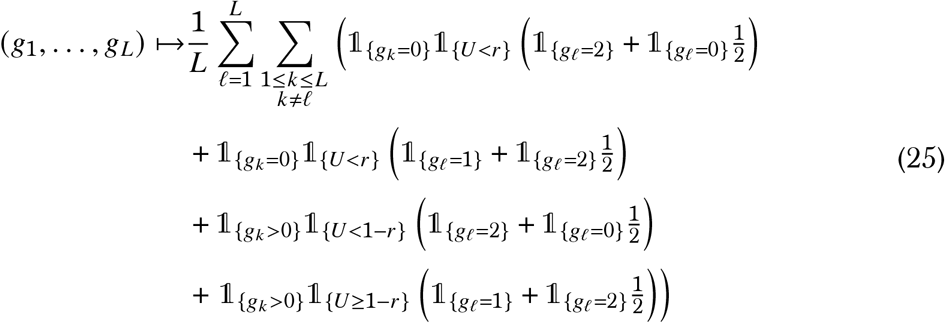

An example for four sites with weight given as in Eq (5) is given in Figure 13.

**Figure 13:**
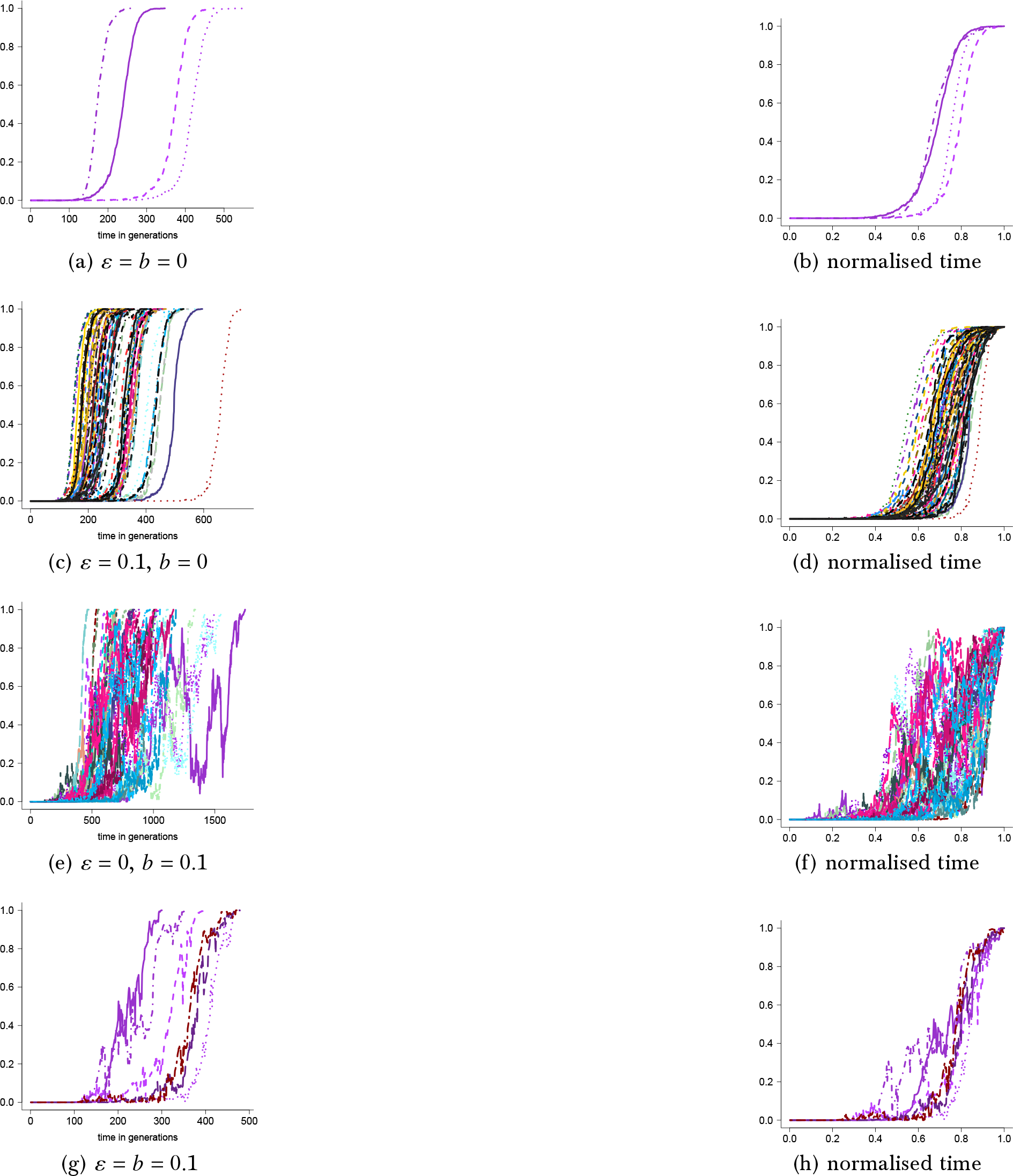
Examples of excursions to fixation for four sites *L* = 4 shown as the number of diploid individuals homozygous for the fit type at all four sites relative to the population size as a function of time with weight computed as in Eq (5) with epistasis among the sites as in Eq (25) for carrying capacity *C* = 10^4^, *α*_1_ = 0.75, *α*_2_ = 3, cutoff *u*(*N*) = *C*, strength of selection (Eq (5)) *s* = 1, bottleneck size *B* = 10^2^ with *ε* and *b* as shown. The time of each excursion in the right panels is normalized by the time to fixation for the excursion. Corresponding numerical estimates of *p* and *τ* are recorded in Table S4. The scale of the abscissa (time axis) may vary between the panels

For a comparison with the additive model in Eq (25) we extend the two-site multiplicative fitness landscape model in Eq (24) to *L* sites to obtain

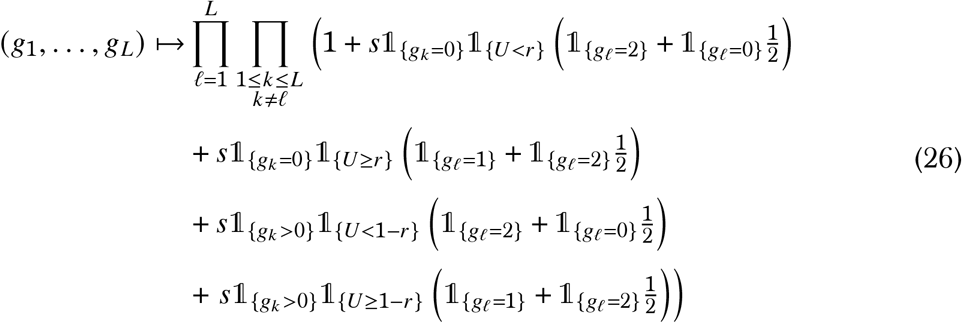

Examples of excursions to fixation at four sites (*L* = 4) with multiplicative fitness as in Eq (26) are recorded in Figure 14 (see Table S4 for numerical estimates of *p* and *τ* corresponding to Figure 13 and Figure 14). Random sweepstakes alone do not alter the fixation trajectories, and only slightly reduce *p* and *τ*. Bottlenecks, however, lead to non-monotonic trajectories where the fit types can spend significant time at intermediate frequencies. Bottlenecks reduce *p* and increase *τ*, while random sweepstakes moderate the effect of bottlenecks on *τ* only, not on *p*.

**Figure 14:**
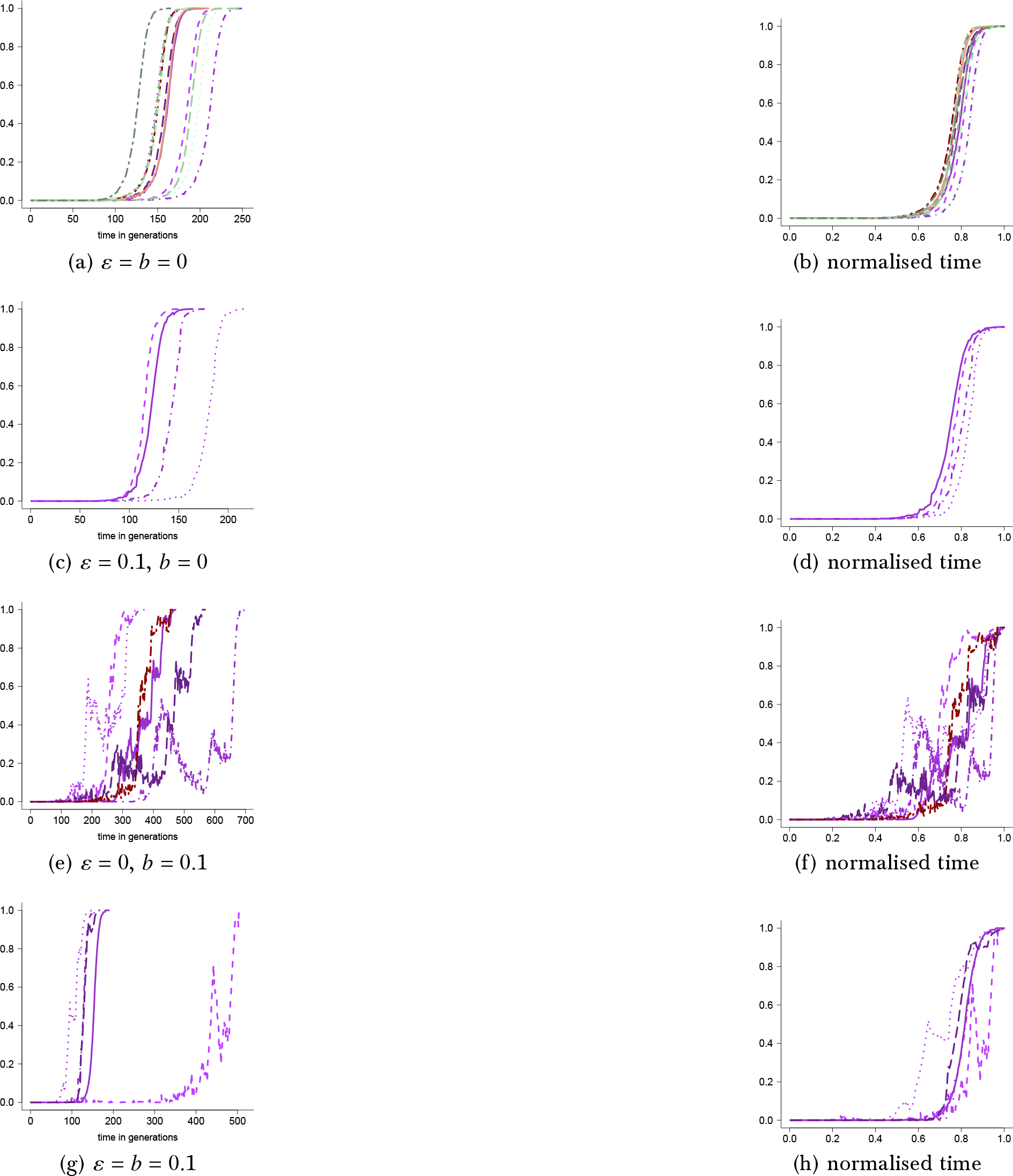
Examples of excursions to fixation for four sites *L* = 4 shown as the number of diploid individuals homozygous for the fit type at all four sites relative to the population size as a function of time with weight given by Eq (26) for for carrying capacity *C* = 10^4^, *α*_1_ = 0.75, *α*_2_ = 3, cutoff *u*(*N*) = *C*, strength of selection *s* = 1, bottleneck size *B* = 10^2^ with *ε* and *b* as shown. The time of each excursion in the right panels is normalized by the time to fixation for the excursion. Corresponding numerical estimates of *p* and *τ* are recorded in Table S4. The scale of the abscissa (time axis) may vary between the panels

## 4 Discussion

In summary, our results show *(i)* fixation of types conferring selective advantage at a number of unlinked sites can occur even when starting with the fit type in a single copy at each site, and despite random sweepstakes and recurrent strong bottlenecks; *(ii)* mixing the dominance mechanisms (i.e. taking the fit type as dominant at one site and recessive at another site) results in trajectories which are mixtures of the trajectories for individual mechanisms; *(iii)* mixtures of over- and underdominance can result in fixation trajectories while restricting to one or the other mechanism does not; *(iv)* random sweepstakes modeled as a mixture of two processes and with an upper bound on the number of potential offspring do not alter the fixation trajectories, but strong recurrent bottlenecks clearly do; *(v)* random sweepstakes and recurrent bottlenecks generally reduce *p*, and strong recurrent bottlenecks generally to a larger extent than random sweepstakes; *(vi)* random sweepstakes generally reduce *τ*, but the effect of recurrent bottlenecks on *τ* depends on the dominance mechanisms.

Fixation of a type conferring selective advantage at two or more unlinked sites has been very little if at all investigated, let alone including further complications such as complex demography or epistasis. Here we provide an algorithm for comparing analytical predictions with simulation results at any number of unlinked sites, and give a few examples of how the stated elements affect selection when acting on at least two unlinked sites. Given the number of parameters involved we have restricted to cases that we think would be of most interest, namely absence of random sweepstakes and bottlenecks (*ε* = *b* = 0), including random sweepstakes but excluding bottlenecks (*ε* = 0.1, *b* = 0), including recurrent bottlenecks and excluding random sweepstakes (*ε* = 0, *b* = 0.1), and including both (*ε* = *b* = 0.1). The restriction to the particular values for *ε* and *b* is only an effort at showing the effects of random sweepstakes and recurrent bottlenecks when either acting ‘strongly’ or not at all. Large families or recurrent bottlenecks can therefore be expected to occur a few times during the time to fixation. The same goes for the bottleneck size (*B*), i.e. we wanted primarily to show the effects of severe bottlenecks; examples of milder bottlenecks can be found in § B. We take *α*_2_ = 3 to guarantee that when *α* = *α*_3_ (see Eq (2)) the probability of large families (at least on the order of the carrying capacity) is so small as to be negligible. It can be shown that when *α* ≥ 2 the ancestral process (tracking the random ancestral relations of sampled gene copies) converges to the Kingman-coalescent(16), and one would expect that, correspondingly, the forward-in-time process tracking a type frequency would converge to the Wright-Fisher diffusion. Throughout we restrict the cutoff (see Eq (2)) *u*(*N*) to the carrying capacity *C*, thereby moderating the effects of random sweepstakes. In the absence of bottlenecks the ancestral process, with *α* ∈ (1, 2), converges to a incomplete Beta-coalescent(16), and when randomizing on *α* as we do here(17). Imposing an upper bound on the number of potential offspring (juveniles) is biologically reasonable; recall that the absence of an upper bound is required to obtain the complete Beta(2 − *α*, *α*)-coalescent(63). Therefore, it is arguably not surprising to see essentially negligible effect of random sweepstakes on the fixation trajectories.

A consistent and pervasive pattern of a convex (U-shaped) site-frequency spectrum is observed in Atlantic cod, and extensive analysis points to a model of recurrent sweeps of strongly advantageous types introduced by mutation(22) as explaining the pattern(4). We claim that our observation that advantageous mutations at unlinked sites can indeed sweep to fixation is further evidence for the conclusion about recurrent sweeps driving the evolution of Atlantic cod. Supposing one observes a convex site-frequency spectrum from a population evolving under recurrent sweeps, i.e. each observed segregating site represents a mutation on it’s way to fixation, can we relate the convex shape to a particular shape of a fixation trajectory? In other words, can the convex shape of a site-frequency spectrum tell us something about the dominance mechanism of the mutations that are sweeping? A convex site-frequency spectrum is, by definition, characterized by a (relatively) high number of low-frequency variants, small number of middle-frequency variants, and a high number of high-frequency variants. If the mutations are recessive then we would most likely not sample them at all since they spend most of the time to fixation in low frequency judging from the fixation trajectories of recessive mutations. A variant that appears as a singleton in a sample is (with high probability) already at a high enough frequency in the population that we have a good chance of picking up at least one copy of it in our sample; the sample frequency of a variant will (with high probability) trail the population frequency. If the mutations are primarily dominant, we would most likely sample them when they are already in high frequency; this might explain the excess of high frequency variants in a convex site-frequency spectrum and the (relative) lack of middle-frequency variants. We also see trajectories with an extended time at high frequency when there is epistasis between two sites with the fit type either dominant or recessive (Figure 1b), or without epistasis but the phenotype is determined by a mixture with the fit type dominant at one site and recessive at another site (Figure 4b or Figure 7b). On the other hand, in the presence of strong recurrent bottlenecks the trajectories tend to spend more time in intermediate frequencies, thus increasing the chance of sampling mutations in that frequency range.

Our algorithm enables us to study how epistasis might affect selection. Through simple models of epistasis we have given a few examples of how epistatic interaction might affect selection. Epistatic interaction between two sites forming a coadapted system has been shown to be important for understanding genetic variation at each site(61). As the molecular mechanisms of epistasis are in general unknown, as well as the degree of epistasis in nature, we certainly do not claim that our models of epistasis are necessarily very realistic. For example we have not considered a potentially important class of epistatic interactions involving diminishing returns epistasis(75). However they do provide further evidence that taking epistatic interactions into account, both in mathematical analysis and in simulations, is necessary for a better understanding of genetic variation in natural populations.

In conclusion, we suggest that dominant advantageous mutations are prevalent contrary to what has been claimed about the dominance mechanism of new mutations(55; 59). The dominance mechanism of new mutations clearly matters for the fate of the mutations, and for the path to fixation when fixation occurs. A mathematical study of the model and its effects on selection might be done using the theory of multitype branching processes (see e.g. (5; 29; 34; 42; 46; 53; 54; 68; 77)). As a somewhat related example, probability of fixation under weak selection can be mathematically investigated using a continuous-state branching process framework(47). However, the model we apply here is quite complicated, involving pairing of diploid individuals, Mendelian inheritance, and potentially complicated epistatic interactions between a number of sites. Nevertheless, studying evolution at more than one site brings us one step towards investigating, in population genetic terms, selection on quantitative traits. Even though quantitative traits have been associated with a large number of QTLs each of small effect(36), QTLs of major effects have been associated with important traits, for example insect resistance in plants(43). Lastly, we have worked under the assumption that epistatic interactions take place between unlinked sites in nature. In principle the algorithm can be extended to include linked sites and recombination, and we propose to leave that for future work. Understanding how the various versions of dominance, and how demography and random sweepstakes shape the excursions to fixation may illuminate how selection, in our case sweeps, may manifest itself in data.

## Acknowledgments

The work was funded by Deutsche Forschungsgemeinschaft (DFG) - Projektnummer 273887127 through DFG SPP 1819: Rapid Evolutionary Adaptation grant STE 325/17 to Wolfgang Stephan; we acknowledge funding by the Icelandic Centre of Research through an Icelandic Research Fund Grant of Excellence no. 185151-051 to Einar Árnason, Katrín Halldórsdóttir, Alison M. Etheridge, Wolfgang Stephan, and BE. BE also acknowledges Start-up module grants through SPP 1819 with Jere Koskela and Maite Wilke Berenguer, and with Iulia Dahmer.

## Code availability

The CWEB (C++) code written for the simulations is available at https://github.com/eldonb/fixation_many_sites

## 5 Supplementary Information

### A Addition to Figure 2

In this section we record an addition to Figure 2 showing excursions to fixation at two sites with weight (5) and weight function given in Eq (10); the only difference to Figure 2 is that here the carrying capacity is *C* = 10^5^. The results are for 10^7^ experiments resulting in eight trajectories to fixation with an average time to fixation 3811.1 generations, 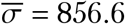 and 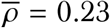. With higher population size we see an even sharper effect of the recessive mechanism, with the excursions spending a great majority of their time with the double homozygotes for the fit type in very low frequency, and by implication with the fit type in low frequency at both sites.

**Figure S1:**
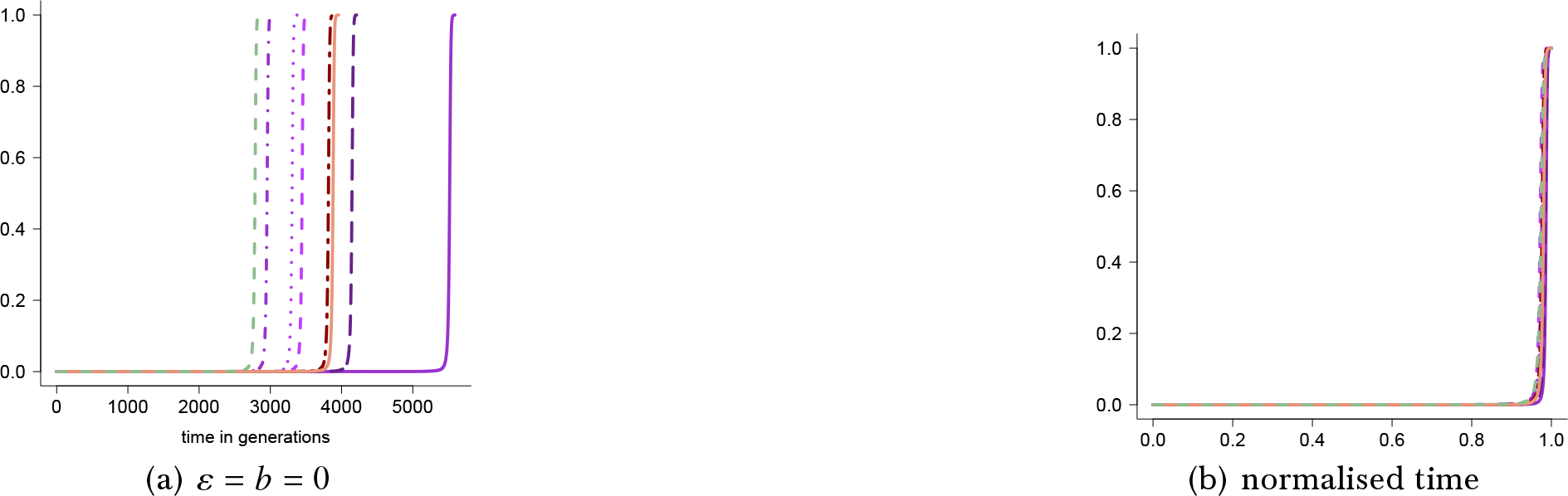
Examples of excursions to fixation of the fit type at two sites shown as the number of individuals homozygous for the fit type at both sites relative to the population size as a function of time, with the weight computed as in Eq (5) with the weight function given in Eq (10) for carrying capacity *C* = 10^5^, *u(N)* = *C*, *α*_1_ = 0.75, *α*_2_ = 3, selection strength (Eq (5)) *s* = 1 with the probability of *α* = *α*_1_ (see Eq (2)) *ε* and the probability of a bottleneck *b* (see Eq (4)) as shown. The scale of the abscissa (time axis) may differ between the graphs

### B Carrying capacity *C* = 10^6^

In this section we record results for two sites when the weight is given by Eq (6), carrying capacity is *C* = 10^6^, and *α*_1_ = 1.05. First we consider the weight function

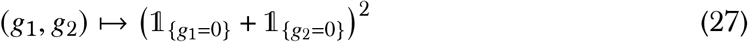

In Eq (27) the fit type is dominant at both sites (see Eq (6)) without epistasis. The results in Figure S2 for the weight function in Eq (27) can be compared to the results in Figure 3.

**Figure S2:**
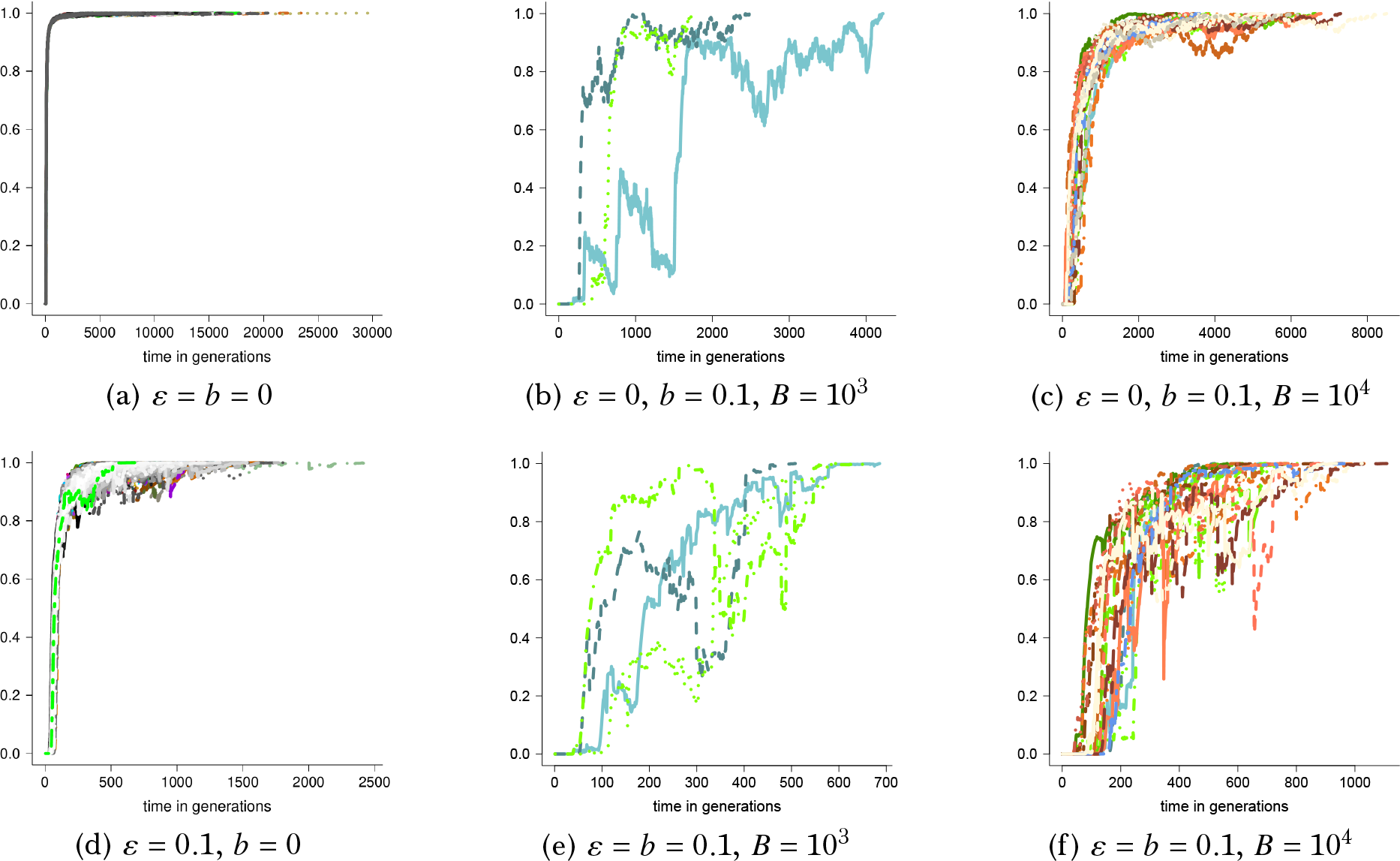
Examples for excursions to fixation of the fit type at two sites shown as the number of individuals homozygous for the fit type at both sites relative to the population size at the time when the fit type is dominant at both sites with weight given by Eq (6) and the weight function given by Eq (27) for carrying capacity *C* = 10^6^, *u(N)* = *C*, *α*_1_ = 1.05, *α*_2_ = 3, selection strength (Eq (6)) *s* = 0.5 with the probability of *α* = *α*_1_ (see Eq (2)) *ε* and the probability of a bottleneck *b* and bottleneck size *B* (see Eq (4)) as shown; the corresponding estimates of *p* (7) and *τ* Eq (8) are recorded in Table S5. The scale of the abscissa (time axis) may differ between the graphs

In Figure S2 the fit type is dominant at both sites (Eq (27)). Now consider the weight function (recall Eq (6))

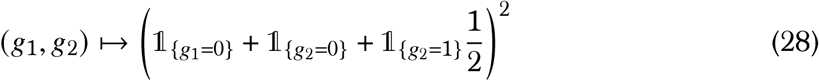

In Eq (28) the fit type is dominant at one site and semidominant at the other site (Figure S3).

**Figure S3:**
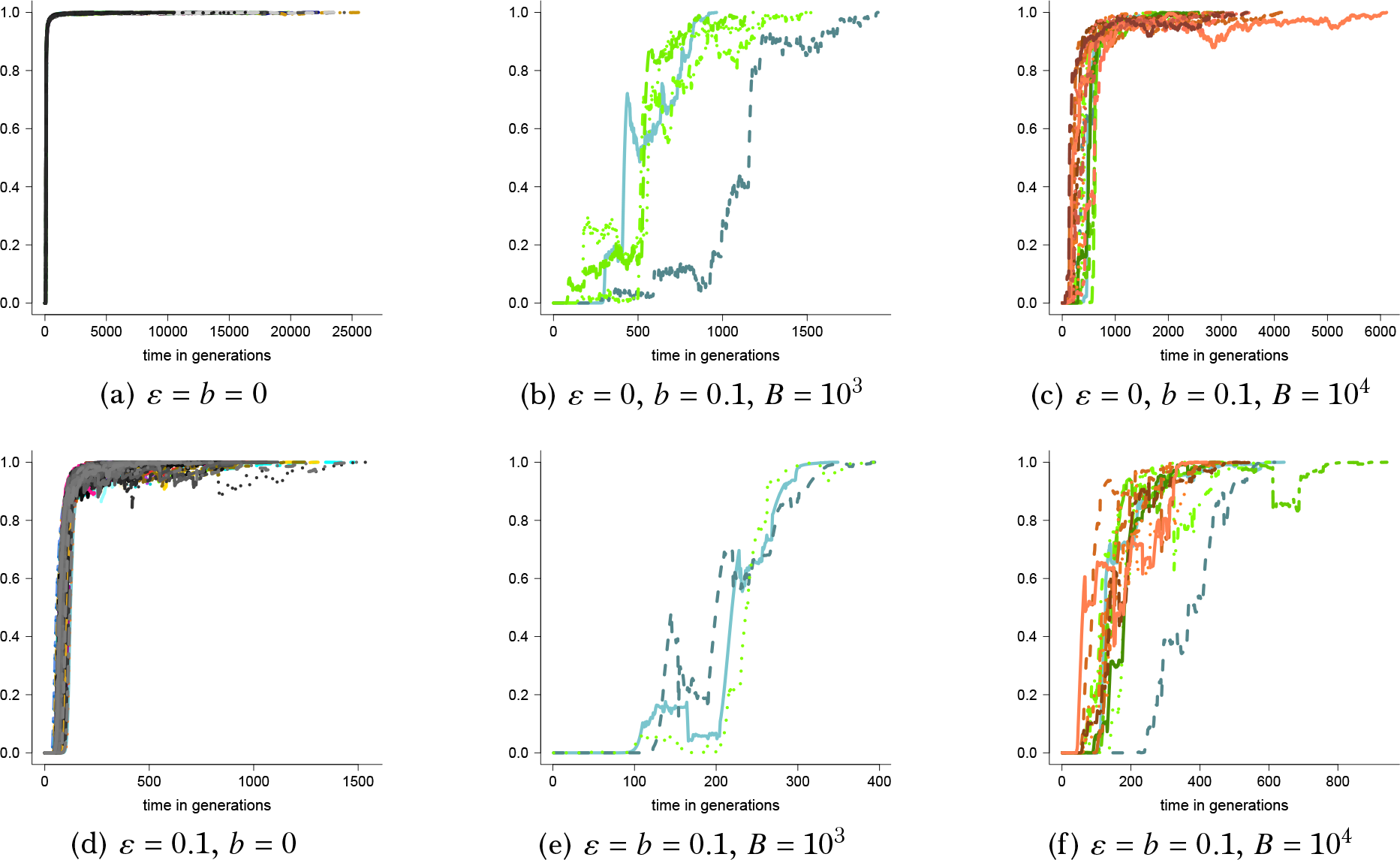
Examples for excursions to fixation shown as the number of individuals at homozygous for the fit type at both sites relative to the population size with the weight function *f* in Eq (6) given by Eq (28) for carrying capacity *C* = 10^6^, *u(N)* = *C*, *α*_1_ = 1.05, *α*_2_ = 3, selection strength (Eq (6)) *s* = 0.5, *ε* and the probability of a bottleneck *b* and bottleneck size *B* (see Eq (4)) as shown; the corresponding estimates of *p* (7) and *τ* Eq (8) are recorded in Table S5. The scale of the abscissa (time axis) may differ between the graphs

Consider the weight function (recall Eq (6))

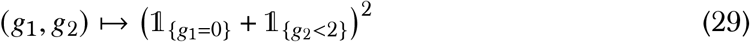

In Eq (29) the fit type is dominant at one site and recessive at the other site without epistasis (see Figure S4). The results in Figure S4 can be compared to Figure 4. Including a recessive site significantly reduces the probability of fixation (Table S5).

**Figure S4:**
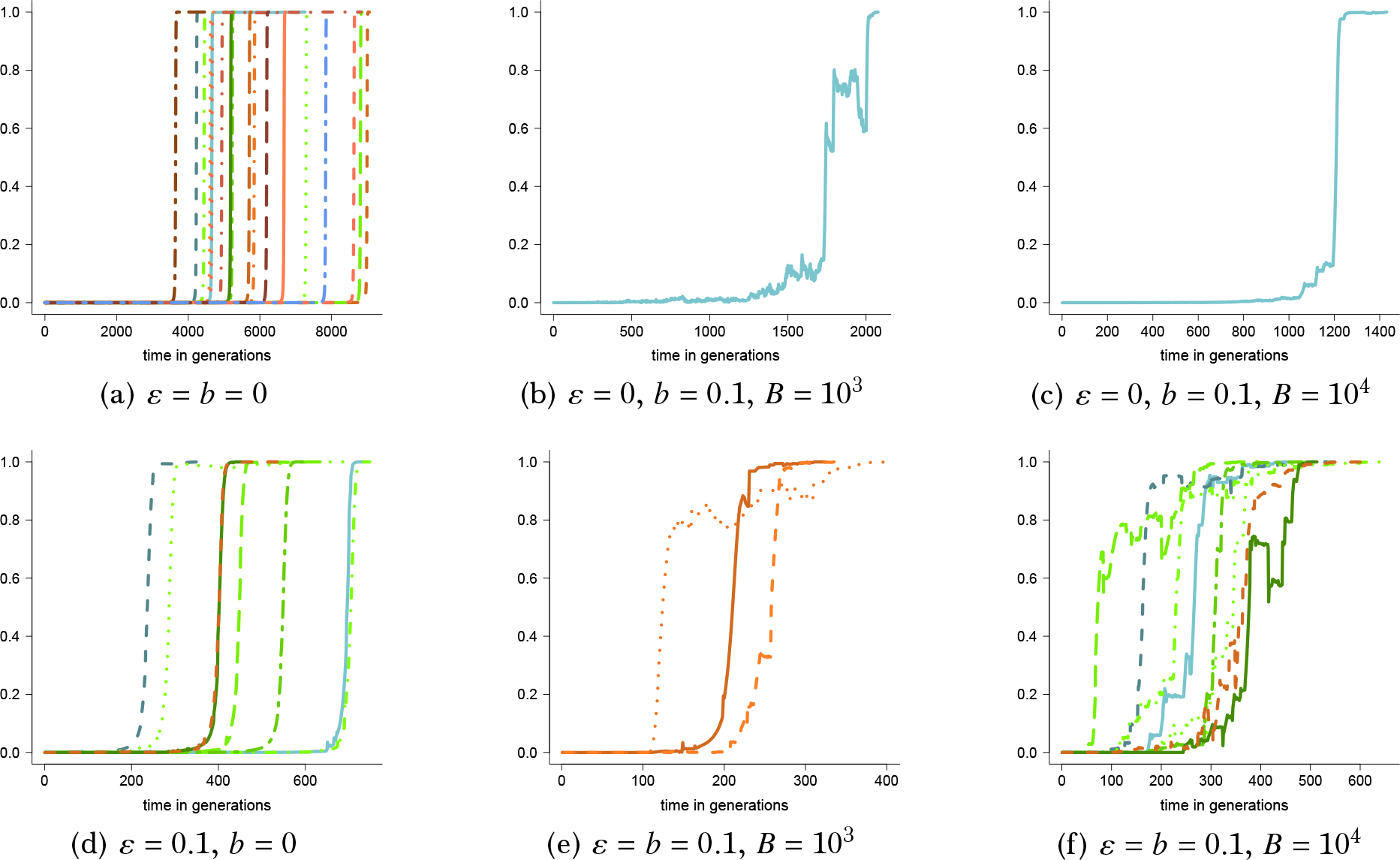
Examples for excursions to fixation shown the number of individuals homozygous for the fit type at both sites relative to the population size as a function of time when the weight is given by Eq (6) and the weight function given by Eq (29) for carrying capacity *C* = 10^6^, *u(N)* = *C*, *α*_1_ = 1.05, *α*_2_ = 3, selection strength (Eq (6)) *s* = 0.5. The corresponding estimates of *p* (7) and *τ* Eq (8) are recorded in Table S5. The scale of the abscissa (time axis) may differ between the graphs

In Figures S2–Fig S4 we considered scenarios where the fit type was dominant at least at one site. Consider the weight function (recall Eq (6))

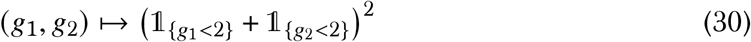

In Eq (30) the fit type is recessive at both sites without epistasis, and the carrying capacity is *C* = 10^5^. The excursions shown in Figure S5 are the results of between 10^6^ and 10^8^ trials.

**Figure S5:**
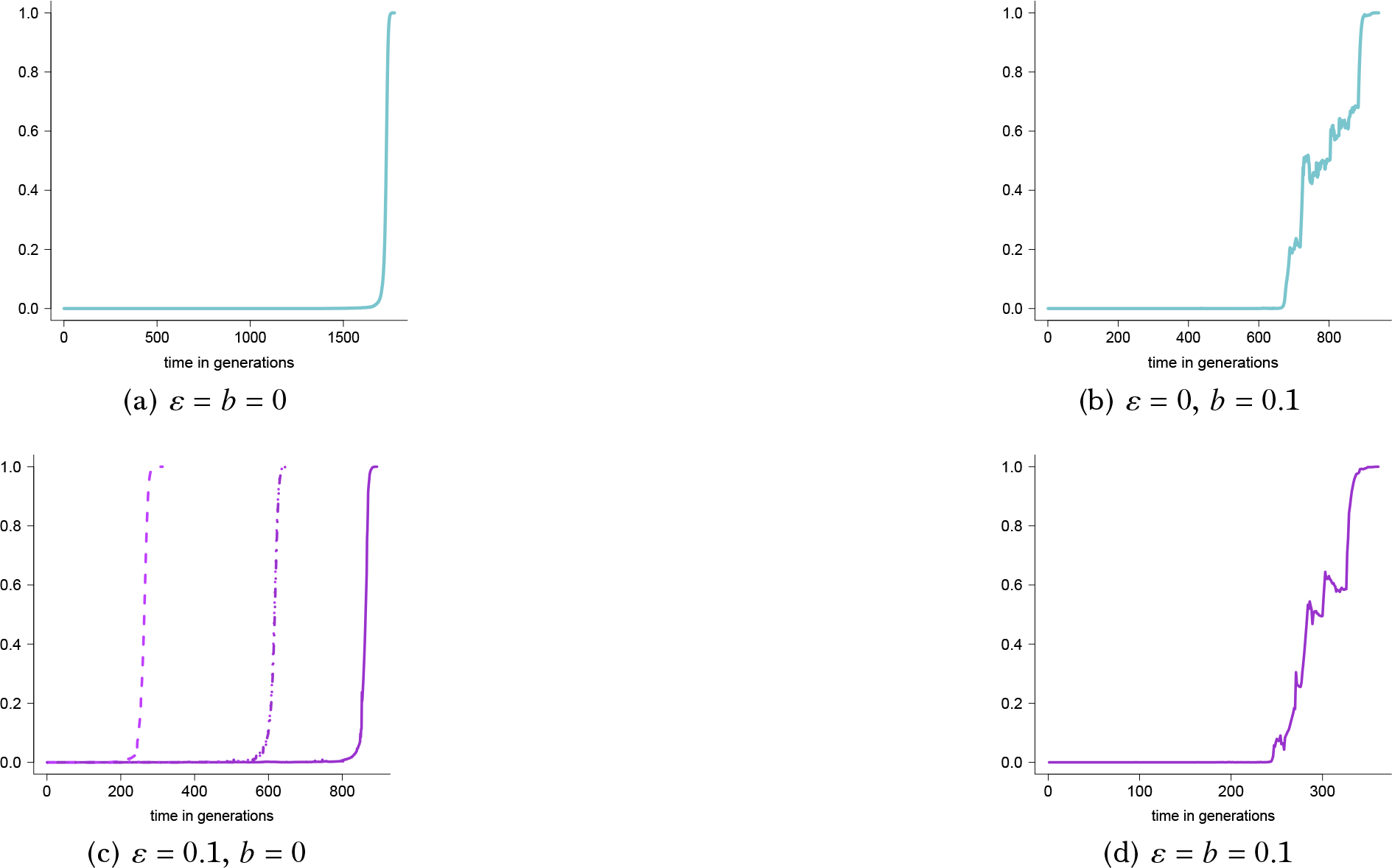
Examples of excursions to fixation shown as *r*_*H*_(*t*) as functions of time *t* where *r*_*H*_(*t*) is the number of individuals at time *t* homozygous for the fit type at both sites relative to the population size at the time for carrying capacity *C* = 10^5^, *α*_1_ = 1.05, *α*_2_ = 3, *u(N)* = *C*, bottleneck size *B* = 10^3^, *ε* and probability of a bottleneck *b* as shown, selection strength (Eq (6)) *s* = 0.5 with the weight function *f* in Eq (6) given as in Eq (30). The scale of the abscissa (time axis) may differ between the graphs

#### B.1 Examples of normalised excursions

In this section we compare excursions to fixation for the fit type at two sites where the time of each excursion is normalised by the time to fixation for the excursion (Figure S6). The fit type is restricted to being either dominant or recessive without epistasis with the weight given as in Eq (6). The weight function is as given in Eq (27) (the fit type dominant at both sites, black lines) and Eq (29) (the fit type dominant at one site and recessive at the other, blue lines).

**Figure S6:**
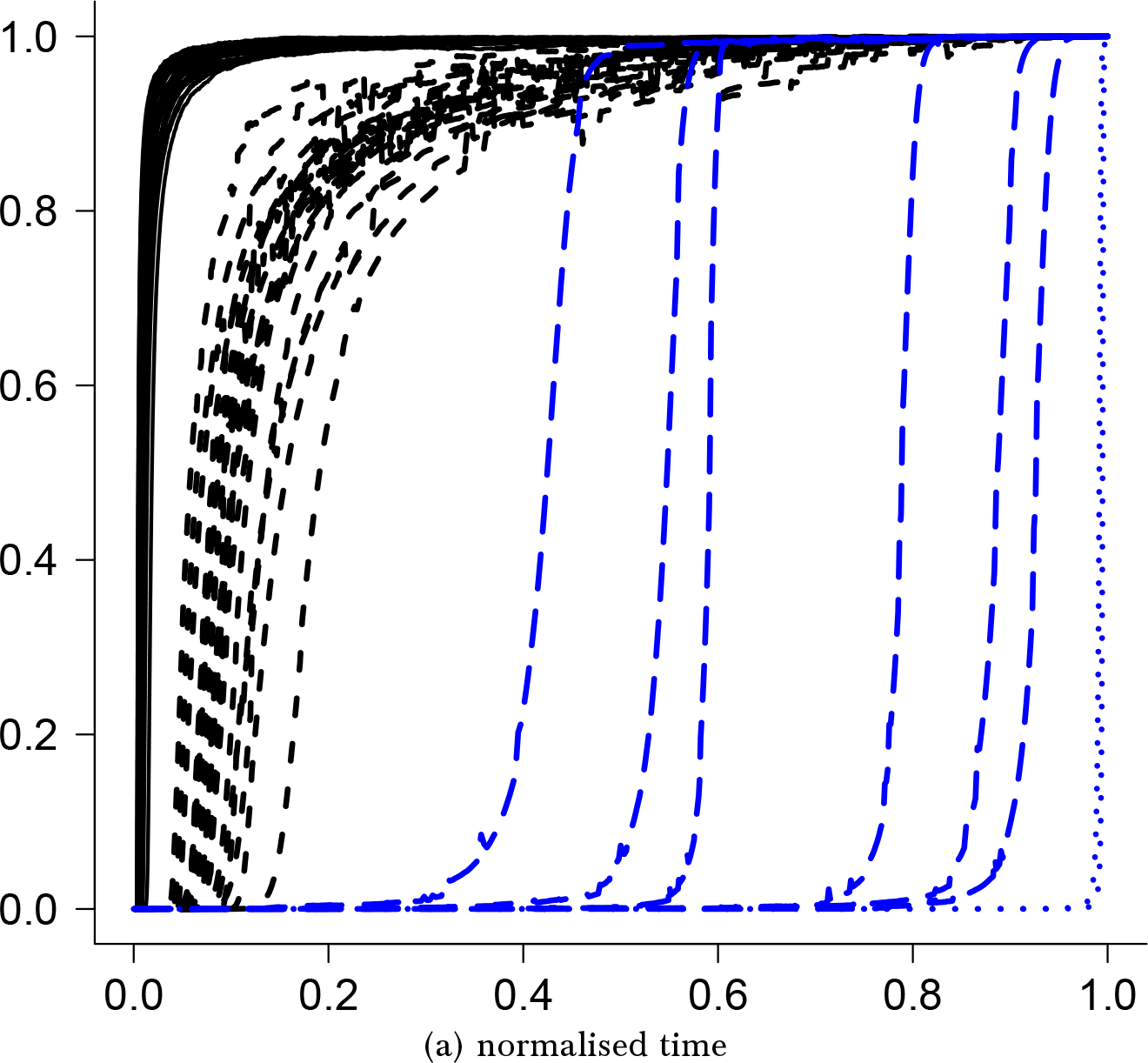
Examples of excursions to fixation for two sites (*L* = 2) shown as number of individuals homozygous for the fit type at both sites relative to the population size as a function of normalised time where the time for each trajectory is normalised by the time to fixation for the trajectory; carrying capacity *C* = 10^6^, *u(N)* = *C*, *α*_1_ = 1.05, *α*_2_ = 3, selection strength *s* = 0.5 and *b* = 0 throughout with the weight function from Eq (6) given as in Eq (27) (black lines) with *ε* = 0 (black solid), *ε* = 0.1 (black dashed), weight function from Eq (6) given as in Eq (29) (blue lines) with *ε* = 0 (blue dotted), *ε* = 0.1 (blue dashed)

#### B.2 Example of epistasis between two sites

In Figure S7 we consider epistasis between two sites with weight computed as in Eq (5) with the weight function given by

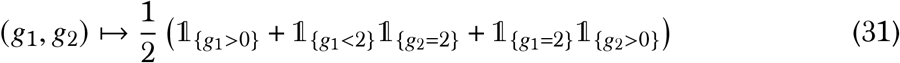

In Eq (31) (compare with Eq (9)) the fit type is dominant at site one, and the dominance mechanism of the fit type at site two depends on the configuration at site one as shown.

**Figure S7:**
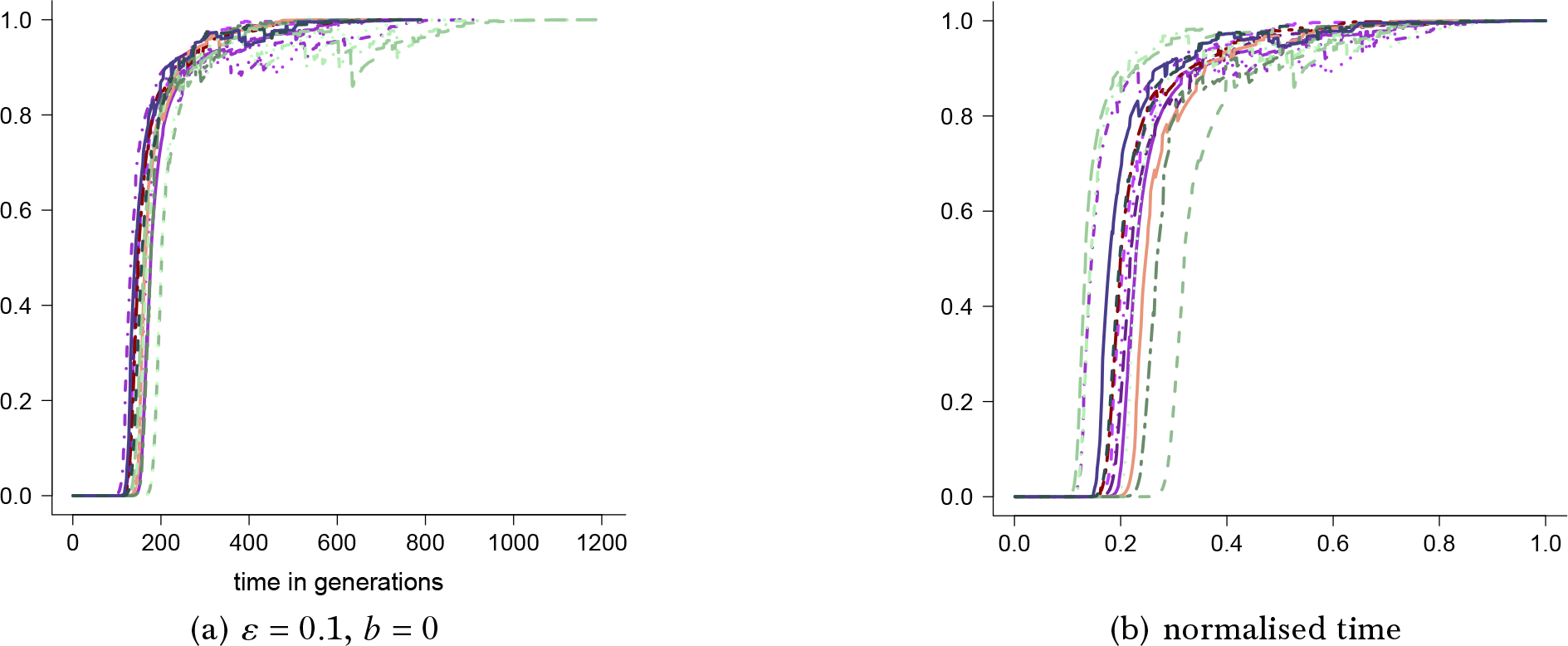
Examples of trajectories to fixation shown as number of individuals homozygous for the fit type at both sites relative to the population size as a function of time with weight given by Eq (5) and weight function as in Eq (31) with carrying capacity *C* = 10^6^, *α*_1_ = 1.05, *ε* = 0.1, cutoff *u(N)* = *C*, *b* = 0, strength of selection *s* = 2; (b) for each excursion in (a) the time to fixation is normalised by the time to fixation for the excursion. The scale of the abscissa (time axis) may vary between panels

### C Further examples

In this section further examples. In Figure S8 for five sites the weight is as given in Eq (6) with weight function

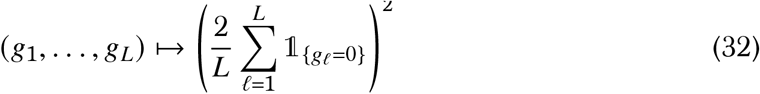

**Figure S8:**
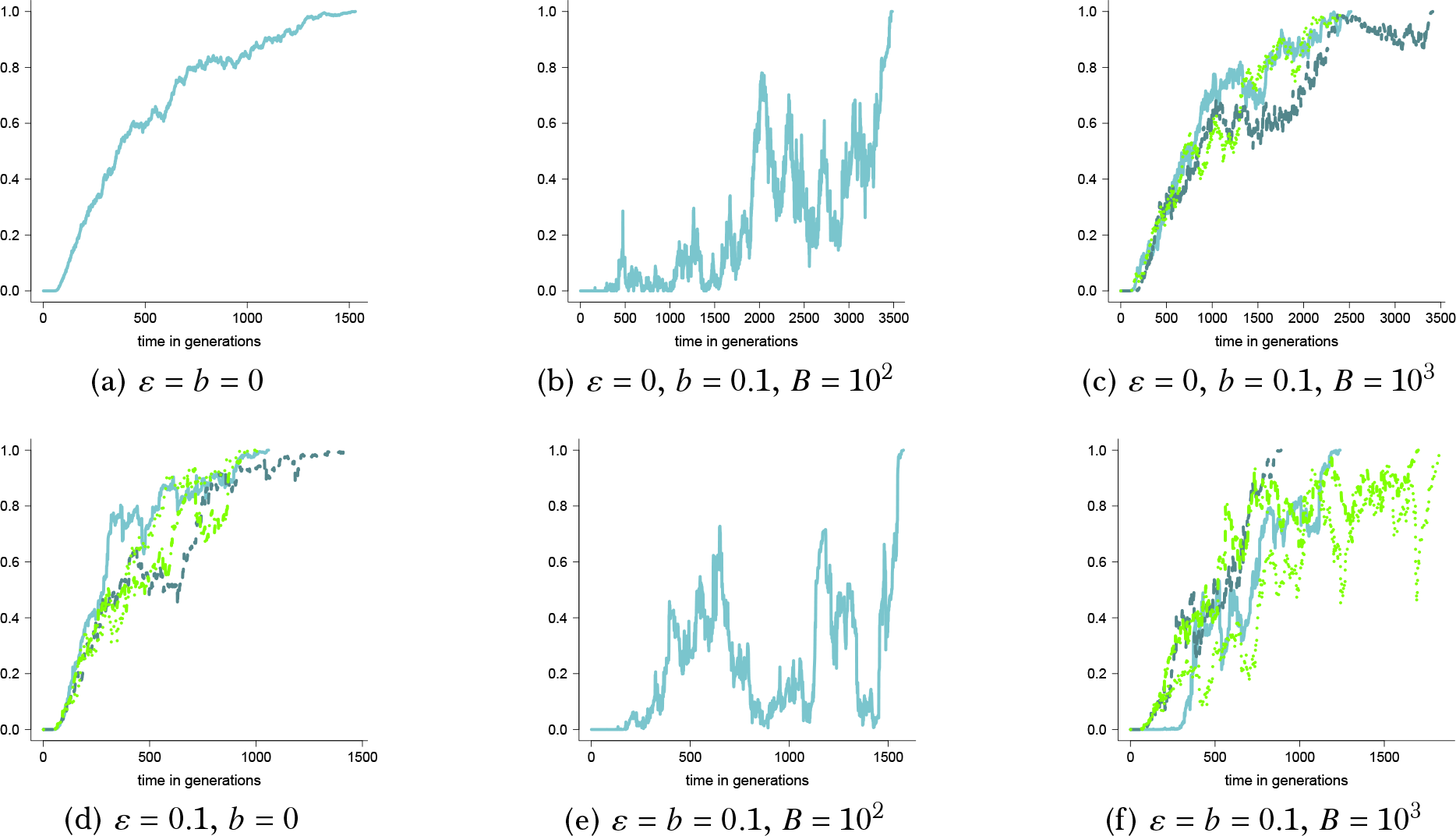
Examples for excursions to fixation shown as the number of individuals at time homozygous for the fit type at all five sites relative to the population size as a function of time with weight as in Eq (6) and weight function as in Eq (32) for carrying capacity *C* = 10^4^, cutoff *u(N)* = *C*, *α*_1_ = 1.05, *α*_2_ = 3, *s* = 0.5, with *ε*, probability of a bottleneck *b*, and bottleneck size *B* as shown. The corresponding estimates of *p* (7) and *τ* Eq (8) are recorded in Table S6. The scale of the abscissa (time axis) may differ between the graphs

In Figure S9 are examples of excursions to fixation for the fit type at two sites with the weight function *f* in Eq (5) given by, with *U* a random uniform on the unit interval,

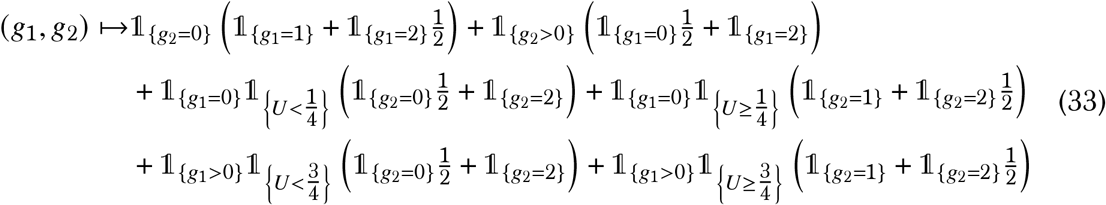

Eq (33) models an additive epistatic fitness landscape model between two sites involving under- 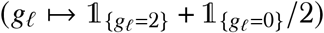 and overdominance 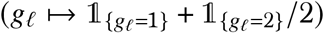 of the fit type, and a random component only for site two. The fit type is overdominant at site one if the type is absent at site one, otherwise the fit type is underdominant at site two. The fit type is underdominant at site two with probability 1/4, otherwise overdominant, if the fit type is absent at site one. If there is at least one copy of the fit type at site one the probabilities of over- and underdominance at site two change as shown is Eq (33).

**Figure S9:**
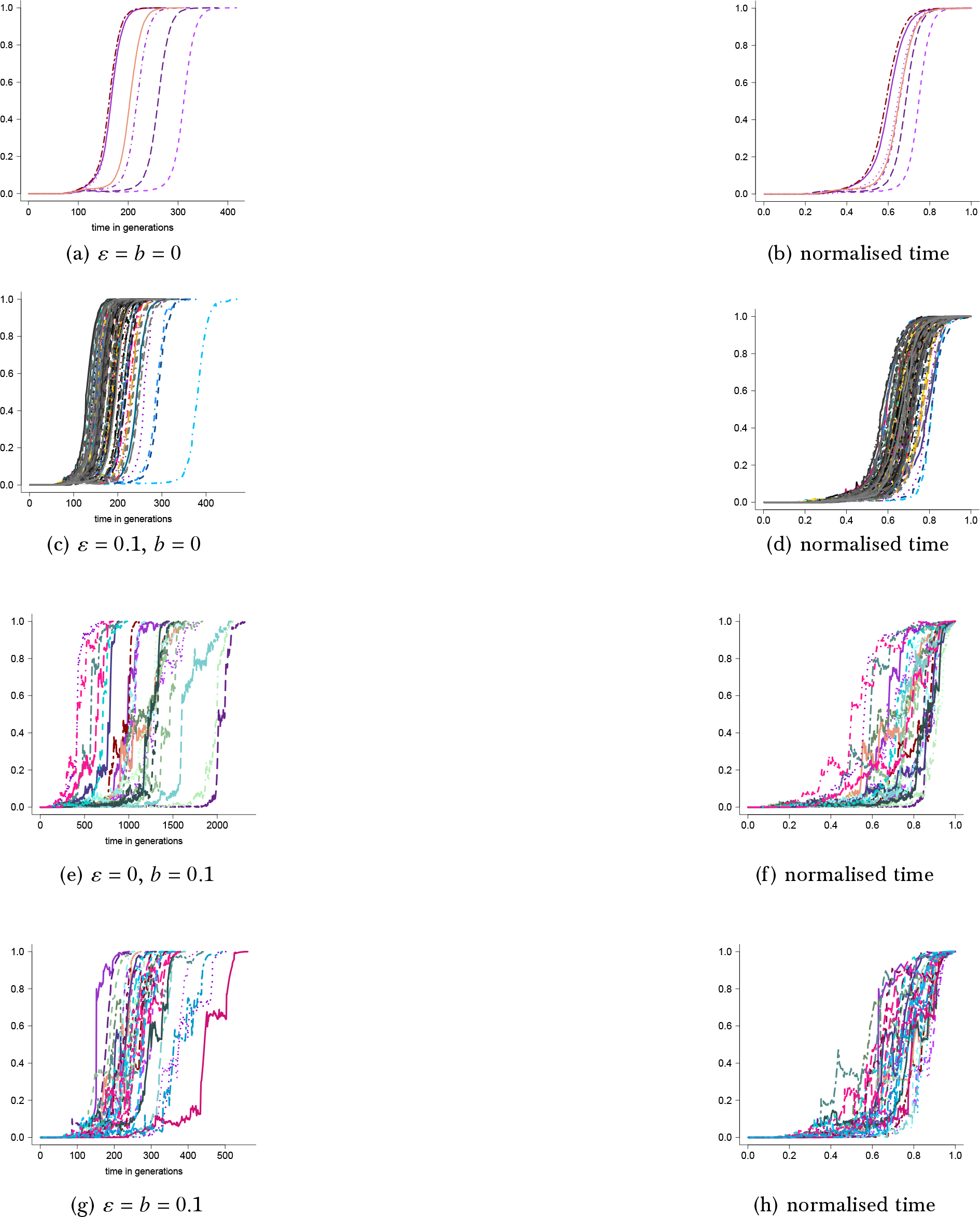
Examples of excursions to fixation shown for two sites shown as the number of diploid individuals homozygous for the fit type at both sites relative to the population size as a function of time with weight in Eq (5) and weight function as in Eq (33) for carrying capacity *C* = 10^5^, *α*_1_ = 0.75, *α*_2_ = 3, *u(N)* = *C*, strength of selection (Eq (5)) *s* = 1, bottleneck size *B* = 10^3^, and *ε* and *b* as shown; the excursions in right panels are with time of each excursion normalized by the time to fixation for the excursion; the corresponding estimates of *p* (Eq (7)) and *τ* (Eq (8)) are recorded in Table S6. The scale of the abscissa (time axis) may vary between panels

### D Numerical estimates for Figures 1–Fig 4

In this section we record numerical estimates of *p* (Eq (7)) and *τ* (Eq (8)) corresponding to Fig 1–Fig 4 (Table S1).

**Table S1:** Numerical estimates of *p* (Eq (7)) and *τ* (Eq (8)) corresponding to Fig 1–Fig 4. Here *m* denotes the number of experiments, *n* the number of experiments resulting in fixation, 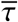 an estimate of *τ*, 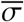 an estimate of the standard deviation of the time to fixation (conditional on fixation), and 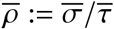.

| case | $m$ | $n$ | $\bar{\tau}$ | $\bar{\sigma}$ | $\bar{\rho}$ |
| --- | --- | --- | --- | --- | --- |
| results corresponding to Figure 1 |  |  |  |  |  |
| $\varepsilon = b = 0$ | $10^6$ | 9 | 1592.1 | 628.6 | 0.39 |
| $\varepsilon = 0.1, b = 0$ | $10^7$ | 34 | 666.1 | 168.4 | 0.25 |
| $\varepsilon = 0, b = 0.1$ | $10^8$ | 12 | 938.6 | 555.1 | 0.59 |
| $\varepsilon = b = 0.1$ | $10^8$ | 13 | 332.9 | 146.2 | 0.44 |
| results corresponding to Figure 2 |  |  |  |  |  |
| $\varepsilon = b = 0$ | $10^6$ | 9 | 997.6 | 242.2 | 0.24 |
| $\varepsilon = 0.1, b = 0$ | $10^7$ | 17 | 390.5 | 131.4 | 0.34 |
| $\varepsilon = 0, b = 0.1$ | $10^8$ | 8 | 626.3 | 306.4 | 0.49 |
| $\varepsilon = b = 0.1$ | $10^8$ | 4 | 403.8 | 205.7 | 0.51 |
| results corresponding to Figure 3 |  |  |  |  |  |
| $\varepsilon = b = 0$ | $10^3$ | 97 | 1351.1 | 472.3 | 0.35 |
| $\varepsilon = 0.1, b = 0$ | $10^3$ | 30 | 566.1 | 192.2 | 0.34 |
| $\varepsilon = 0, b = 0.1$ | $10^4$ | 11 | 915.0 | 314.4 | 0.34 |
| $\varepsilon = b = 0.1$ | $10^4$ | 5 | 291.2 | 117.6 | 0.40 |
| results corresponding to Figure 4 |  |  |  |  |  |
| $\varepsilon = b = 0$ | $10^4$ | 7 | 1214.9 | 570.4 | 0.47 |
| $\varepsilon = 0.1, b = 0$ | $10^5$ | 19 | 487.9 | 151.8 | 0.31 |
| $\varepsilon = 0, b = 0.1$ | $10^7$ | 19 | 663.3 | 184.3 | 0.28 |
| $\varepsilon = b = 0.1$ | $10^7$ | 23 | 311.1 | 97.4 | 0.31 |

### E Numerical estimates for Figures 5–Fig 8

In this section we record numerical estimates of *p* (Eq (7)) and *τ* (Eq (8)) corresponding to Figures 5–Fig 8 (Table S2).

**Table S2:** Estimates of *p* (Eq (7)) and *τ* (Eq (8)) corresponding to Figures 5–Fig 8. Here *m* denotes the number of experiments, *n* the number of experiments resulting in fixation, 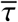 an estimate of *τ*, 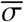 an estimate of the standard deviation of the time to fixation (conditional on fixation), and 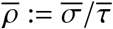.

| case | $m$ | $n$ | $\bar{\tau}$ | $\bar{\sigma}$ | $\bar{\rho}$ |
| --- | --- | --- | --- | --- | --- |
| results corresponding to Figure 5 |  |  |  |  |  |
| $\varepsilon = b = 0$ | $10^2$ | 4 | 483.8 | 377.5 | 0.78 |
| $\varepsilon = 0.1, b = 0$ | $10^3$ | 13 | 196.8 | 112.9 | 0.57 |
| $\varepsilon = 0, b = 0.1$ | $10^4$ | 3 | 359.7 | 66.0 | 0.18 |
| $\varepsilon = b = 0.1$ | $10^4$ | 6 | 196.0 | 39.7 | 0.20 |
| results corresponding to Figure 6 |  |  |  |  |  |
| $\varepsilon = b = 0$ | $10^2$ | 11 | 1326.9 | 685.8 | 0.52 |
| $\varepsilon = 0.1, b = 0$ | $10^2$ | 3 | 476.3 | 210.3 | 0.44 |
| $\varepsilon = 0, b = 0.1$ | $10^4$ | 10 | 644.3 | 324.4 | 0.50 |
| $\varepsilon = b = 0.1$ | $10^4$ | 6 | 305.0 | 68.6 | 0.22 |
| results corresponding to Figure 7 |  |  |  |  |  |
| $\varepsilon = b = 0$ | $10^4$ | 8 | 1102.1 | 467.6 | 0.42 |
| $\varepsilon = 0.1, b = 0$ | $10^4$ | 2 | 566.0 | 246.1 | 0.43 |
| $\varepsilon = 0, b = 0.1$ | $10^6$ | 2 | 555.0 | 155.6 | 0.28 |
| $\varepsilon = b = 0.1$ | $10^5$ | 2 | 255.0 | 26.9 | 0.11 |
| results corresponding to Figure 8 |  |  |  |  |  |
| $\varepsilon = b = 0$ | $10^6$ | 12 | 734.8 | 170.0 | 0.23 |
| $\varepsilon = 0.1, b = 0$ | $10^6$ | 4 | 345.0 | 126.7 | 0.37 |
| $\varepsilon = 0, b = 0.1$ | $10^8$ | 5 | 297.0 | 89.8 | 0.30 |
| $\varepsilon = b = 0.1$ | $10^8$ | 12 | 185.6 | 76.8 | 0.41 |

### F Numerical estimates corresponding to Figure 10– Figure 11

The numerical estimates of *p* (Eq (7)) and *τ* (Eq (8)) corresponding to Figure 10 and Figure 11 are recorded in Table S3.

### G Numerical estimates corresponding to Figure 13 and Figure 14

In this section record numerical estimates of *p* (Eq (7)) and *τ* (Eq (8)) corresponding to Figure 13 and Figure 14 (Table S4).

**Table S3:** Estimates of *p* (Eq (7)) and *τ* (Eq (8)) corresponding to Figure 10 and Figure 11. Here *m* denotes the number of experiments, *n* the number of experiments resulting in fixation, 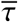 an estimate of *τ*, 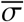 an estimate of the standard deviation of the time to fixation (conditional on fixation), and 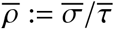.

| case | $m$ | $n$ | $\bar{\tau}$ | $\bar{\sigma}$ | $\bar{\rho}$ |
| --- | --- | --- | --- | --- | --- |
| results corresponding to Figure 10 |  |  |  |  |  |
| $\varepsilon = b = 0$ | $10^4$ | 36 | 439.3 | 173.5 | 0.39 |
| $\varepsilon = 0.1, b = 0$ | $10^3$ | 7 | 361.1 | 125.9 | 0.35 |
| $\varepsilon = 0, b = 0.1$ | $10^5$ | 9 | 803.8 | 539.9 | 0.67 |
| $\varepsilon = b = 0.1$ | $10^5$ | 5 | 355.8 | 126.7 | 0.36 |
| results corresponding to Figure 11 |  |  |  |  |  |
| $\varepsilon = b = 0$ | $10^3$ | 30 | 342.1 | 158.8 | 0.46 |
| $\varepsilon = 0.1, b = 0$ | $10^3$ | 7 | 271.1 | 80.2 | 0.30 |
| $\varepsilon = 0, b = 0.1$ | $10^5$ | 8 | 527.6 | 228.3 | 0.43 |
| $\varepsilon = b = 0.1$ | $10^5$ | 7 | 329.0 | 103.2 | 0.31 |

**Table S4:** Numerical estimates of *p* (Eq (7)) and *τ* (Eq (8)) for Figure 13 and Figure 14; *m* is the number of experiments, *n* the number of excursions resulting in fixation, 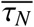 is the average number of generations to fixation (conditional on fixation), 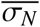 is the standard deviation of the time to fixation (conditional on fixation), and 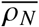 the coefficient of variation of the time to fixation.

| case | $m$ | $n$ | $\overline{\tau_N}$ | $\overline{\sigma_N}$ | $\overline{\rho_N}$ |
| --- | --- | --- | --- | --- | --- |
| results corresponding to Figure 13 |  |  |  |  |  |
| $\varepsilon = b = 0$ | $10^3$ | 4 | 407.5 | 131.3 | 0.32 |
| $\varepsilon = 0.1, b = 0$ | $10^5$ | 75 | 371.8 | 95.5 | 0.26 |
| $\varepsilon = 0, b = 0.1$ | $10^7$ | 30 | 971.0 | 294.7 | 0.30 |
| $\varepsilon = b = 0.1$ | $10^7$ | 6 | 415.2 | 76.8 | 0.19 |
| results corresponding to Figure 14 |  |  |  |  |  |
| $\varepsilon = b = 0$ | $10^3$ | 12 | 209.8 | 26.5 | 0.13 |
| $\varepsilon = 0.1, b = 0$ | $10^3$ | 4 | 176.8 | 29.3 | 0.17 |
| $\varepsilon = 0, b = 0.1$ | $10^5$ | 6 | 486.0 | 131.8 | 0.27 |
| $\varepsilon = b = 0.1$ | $10^5$ | 5 | 234.8 | 159.0 | 0.68 |

### H Numerical estimates corresponding to Fig S2–Fig S4

In this section we record the numerical estimates of *p* (Eq (7)) and *τ* (Eq (8)) corresponding to Figures S2–Fig S4 (Table S5).

**Table S5:** Numerical estimates of *p* (Eq (7)) and *τ* (Eq (8)) corresponding to Fig S2– Fig S4; *m* is the number of experiments, *n* the number of excursions resulting in fixation, 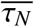 is the average number of generations to fixation (conditional on fixation), 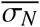 is the standard deviation of the time to fixation (conditional on fixation), and 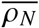 the coefficient of variation of the time to fixation.

| case | $m$ | $n$ | $\overline{\tau_N}$ | $\overline{\sigma_N}$ | $\overline{\rho_N}$ |
| --- | --- | --- | --- | --- | --- |
| results corresponding to Figure S2 |  |  |  |  |  |
| $\varepsilon = b = 0$ | $10^3$ | 253 | 9949.9 | 3831.9 | 0.39 |
| $\varepsilon = 0, b = 0.1, B = 10^3$ | $10^3$ | 3 | 2835.3 | 1254.4 | 0.44 |
| $\varepsilon = 0, b = 0.1, B = 10^4$ | $10^3$ | 22 | 4752.8 | 1698.7 | 0.36 |
| $\varepsilon = 0.1, b = 0$ | $10^3$ | 210 | 897.7 | 286.0 | 0.32 |
| $\varepsilon = b = 0.1, B = 10^3$ | $10^3$ | 4 | 614.8 | 79.0 | 0.13 |
| $\varepsilon = b = 0.1, B = 10^4$ | $10^3$ | 20 | 797.0 | 169.6 | 0.21 |
| results corresponding to Figure S3 |  |  |  |  |  |
| $\varepsilon = b = 0$ | $10^3$ | 233 | 8201.6 | 4031.4 | 0.49 |
| $\varepsilon = 0, b = 0.1, B = 10^3$ | $10^3$ | 5 | 1421.0 | 365.6 | 0.26 |
| $\varepsilon = 0, b = 0.1, B = 10^4$ | $10^3$ | 17 | 3033.5 | 1059.9 | 0.35 |
| $\varepsilon = 0.1, b = 0$ | $10^3$ | 158 | 698.7 | 232.4 | 0.33 |
| $\varepsilon = b = 0.1, B = 10^3$ | $10^3$ | 3 | 380.7 | 27.5 | 0.07 |
| $\varepsilon = b = 0.1, B = 10^4$ | $10^3$ | 13 | 541.3 | 148.6 | 0.27 |
| results corresponding to Figure S4 |  |  |  |  |  |
| $\varepsilon = b = 0$ | $10^5$ | 18 | 6466.2 | 1659.9 | 0.26 |
| $\varepsilon = 0, b = 0.1, B = 10^3$ | $10^5$ | 1 | 2077.0 | – | – |
| $\varepsilon = 0, b = 0.1, B = 10^4$ | $10^6$ | 1 | 1431.0 | – | – |
| $\varepsilon = 0.1, b = 0$ | $10^6$ | 8 | 606.8 | 132.7 | 0.22 |
| $\varepsilon = b = 0.1, B = 10^3$ | $10^7$ | 3 | 357.7 | 39.1 | 0.11 |
| $\varepsilon = b = 0.1, B = 10^4$ | $10^7$ | 8 | 529.9 | 85.3 | 0.16 |

### I Numerical estimates corresponding to Fig S8 and Fig S9

In this section we record numerical estimates of *p* (Eq (7)) and *τ* (Eq (8)) corresponding to Figure S8 and Figure S9 (Table S6).

**Table S6:**
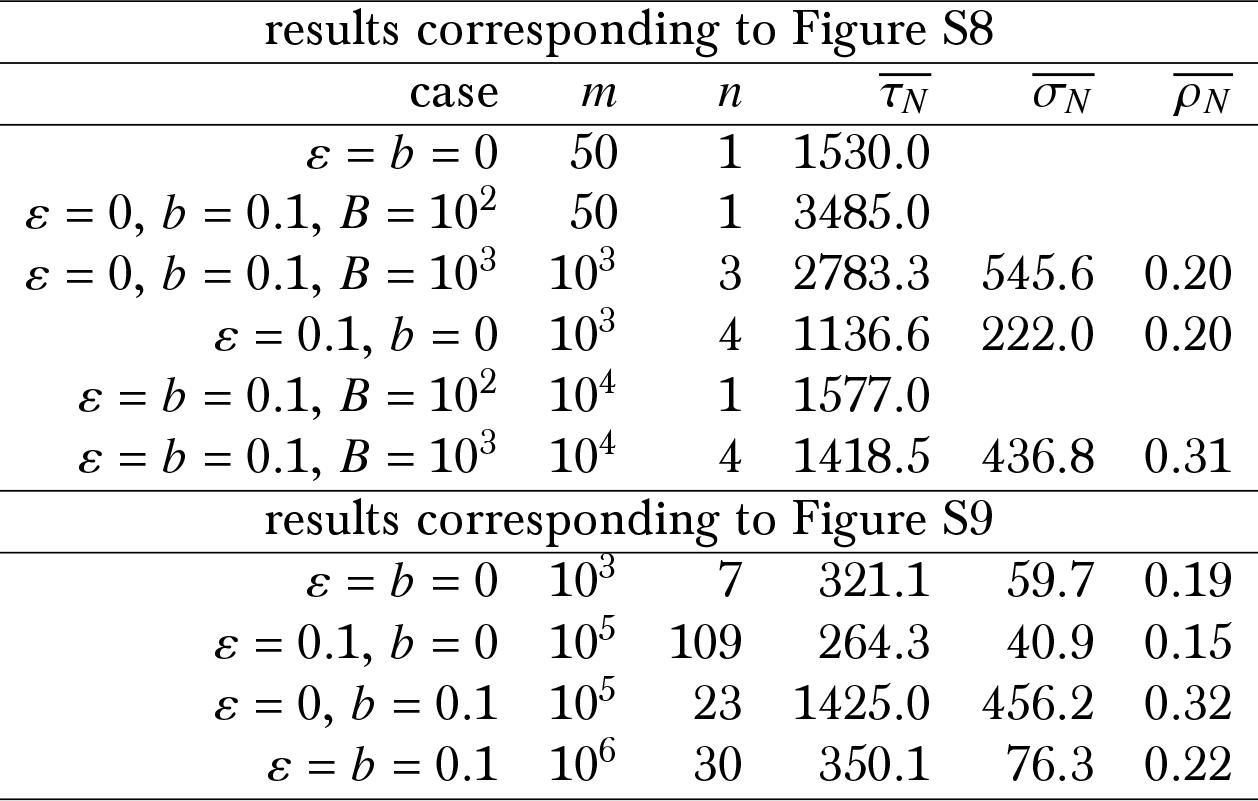
Numerical estimates of *p* (Eq (7)) and *τ* (Eq (8)) corresponding to Figure S8 and Figure S9; *m* is the number of experiments, *n* the number of excursions resulting in fixation, 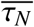 is the average number of generations to fixation (conditional on fixation), 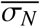 is the standard deviation of the time to fixation (conditional on fixation), and 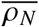 the coefficient of variation of the time to fixation.

### J Dominance mechanisms

In this section we give a brief overview of dominance mechanisms (Table S7). Writing *z*_0_, *z*_1_, *z*_2_ for the phenotypic value of the homozygote for the wild type resp. the heterozygote resp. the homozygote for the fit type, Table S7 shows the relation between them for each dominance mechanism assuming fitness increases with the phenotypic value.

**Table S7:** Main dominance mechanisms.

| term | relation |
| --- | --- |
| overdominance | $z_0 < z_2 < z_1$ |
| dominance | $z_0 < z_1 = z_2$ |
| semidominance | $z_0 < z_1 < z_2$ |
| recessive | $z_0 = z_1 < z_2$ |
| underdominance | $z_1 < z_0 < z_2$ |

### K pseudocode

In this section we formulate the algorithm in a pseudocode. Let *Y(t)* ≔ (*Y*_1_(*t*),…, *Y*_*L*_(*t*)) ϵ [0,1]^*L*^ be the L-site type frequency process, where *Y*_ℓ_(*t*) is the number of copies of the type conferring selective advantage at site *ℓ* relative to the population size at time *t*. Write *n* ≔ {1,2,…, *n*} for *n* ϵ ℕ, *N*_*t*_ for the population size at time *t*, *B* for the population size right after a bottleneck, *p* for the probability of a bottleneck in a given generation, *g* = (*g*_1_,…, *g*_*L*_) an L-site type, and *f* (*g*) a function for how *g* determines the trait value

1. 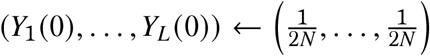 with population set in the starting configuration as described above
2. while({Π_*ℓ*∈[*L*]_ *Y*_ℓ_ (*t*) > 0} ⋂ {Σ_*ℓ*∈[*L*]_ *Y*_ℓ_ (*t*) < *L*) then for generation *t* + 1 started with *N*_*t*_ individuals:
  a. *U* ← random uniform on [0, 1]
  b. 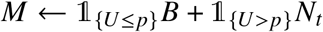 where *p* is the probability of a bottleneck
  c. the *M* individuals in the case of a bottleneck are sampled uniformly at random and without replacement from the *N*_*t*_ individuals
  d. sample *X*_1_,…, *X*_⌊*M*/2⌋_ random number of juveniles produced independently by ⌊*M*/2⌋ parent pairs and assign types according to Mendel’s laws
  e. *S*_⌊*M*/2⌋_ ← *X*_1_ + … + *X*_⌊*M*/2⌋_ the total number of juveniles
  f. sample a random exponential *E* ← Exp exp (−*sf*(*g*))) for each juvenile with rate exp (−*sf*(*g*))
  g. 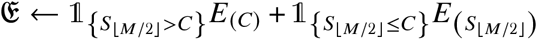 where *E*_(*j*)_ is the *j* th smallest exponential out of the *S*_⌊*M*/2⌋_ exponentials from (d)
  h. a juvenile with sampled exponential *E* survives if 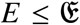
3. record the result of the trajectory according to {Π_*ℓ*∈[*L*]_ *Y*_ℓ_ (*t*) = 0} or {Σ_*ℓ*∈[*L*]_ *Y*_ℓ_ (*t*) = *L*}

### L Mendelian probabilities

Given *L*-site types 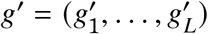 and 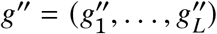 of parents, the probability that an (random) offspring type *G* is of type *g* = (*g*_1_,…, *g*_*L*_) is

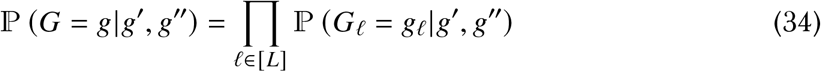

since the gene copies at each site segregate independently and recalling that 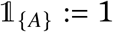 if *A* holds, and zero otherwise,

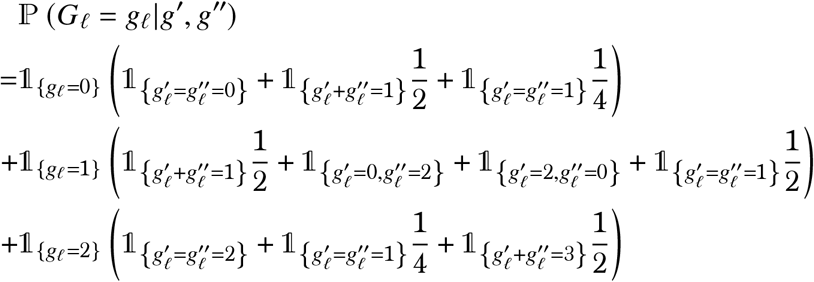

where 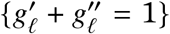 denotes the event that one parent is of type 0 (homozygote for the wild type) and the other parent a heterozygote (type 1), and 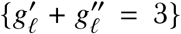 the event that one parent is a heterozygote and the other parent homozygous for the fit type (type 2). Writing *r* ≔ ℙ (*G* = *g*|*h*′, *h*″), the probability that a pair of parents of type {*h*′, *h*″} produces *y* juveniles of type *g* is then

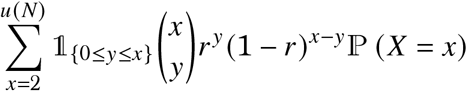

where ℙ (*X* = *x*) is the probability of *X* juveniles according to Eq (2). Since any given type can be produced from different parent types, the total number of juveniles of a particular type produced in any given generation is obtained by adding up the contributions from all the parent pairs. The random number of juveniles of each type produced by a parent pair with given types, conditional on the total number of juveniles *X* produced by the pair, is a multinomial with parameters *X* and the probabilities in Eq (34).

Let *ν* ≔ (*ν*_1_*,…, *ν*_*N*_*) denote the number of surviving offspring derived from the current *N* individuals. In our model with the production of juveniles the probability of an offspring vector *v* taking values (*v*_1_,…,*v*_*N*_) ϵ {0,1,…, *N*}^*N*^ with 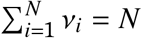 in a haploid population of fixed size *N* is then (assuming *X*_1_ + … + *X*_*N*_ ≥ *N* almost surely)

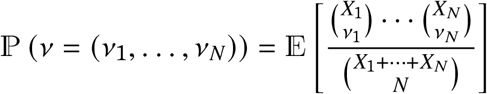

where (*X*_1_,…, *X*_*N*_) denotes the random number of juveniles produced according to Eq (2) (adapted to a haploid population). Supposing there are two types, and we denote by {*Y*_*t*_} the Markov chain tracking the number of one of the types, then in a constant haploid population of size *N* in the absence of selection exchangeability gives

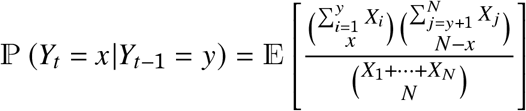

